# Structure-specific Mini-Prion Model for Alzheimer’s Disease Tau Fibrils

**DOI:** 10.1101/2025.06.23.661111

**Authors:** Vishnu Vijayan, Gregory E. Merz, Karen Tsay, Andrew P. Longhini, Samuel Lobo, Athena Quddus, Kristi Lynn S. Nakagawa, Michael P. Vigers, Arthur A. Melo, Eric Tse, Joan-Emma Shea, M. Scott Shell, Kenneth S. Kosik, Daniel R. Southworth, Songi Han

## Abstract

A critical discovery of the past decade is that tau protein fibrils adopt disease-specific hallmark structures in each tauopathy. The faithful generation of synthetic fibrils adopting hallmark structures that can serve as targets for developing diagnostic and/or therapeutic strategies remains a grand challenge. We report on a rational design of synthetic fibrils built of a short peptide that adopts a critical structural motif in tauopathy fibrils found in Alzheimer’s Disease (AD) and Chronic Traumatic Encephalopathy (CTE). They serve as minimal prions with exquisite seeding competency, *in vitro* and in tau biosensor cells, for recruiting tau constructs ten times larger its size *en route* to AD or CTE fibril structures. We demonstrate that the generation of AD and CTE-like fibril structures is dramatically catalyzed in the presence of mini-AD prions and further influenced by salt composition in solution. Double Electron-Electron Resonance studies confirmed the preservation of AD-like folds across multi-generational seeding. Fibrils formed with the full AD/CTE-like core show strong seeding competency, with their templating effect dominating over the choice of salt composition that tunes the initial selection of AD- and CTE-like fibril populations. The mini-AD prions serve as a potent catalyst with templating capabilities that offer a novel strategy to design pathological tau fibril models.

## Introduction

Recent advances in single particle cryogenic electron microscopy (cryo-EM) have profoundly enhanced our understanding of tauopathies, a group of neurodegenerative diseases characterized by the deposition of fibrillar aggregates of the microtubule-associated protein tau (MAPT) in the brain. Among the most significant revelations is that the brain deposits made of tau fibrils adopt distinct structures unique to each tauopathy across different patient brains from the same disease (1). The genesis and mechanisms driving the highly selective shape propagation underlying the tau folds in each disease-specific tau fibril remain elusive. A prevalent hypothesis that guides this study is that structural selection is key to pathological tau fibril propagation and is driven by a ‘prion-like’ disease progression, in which an elongating fibril end surface made of unique monolayer folds of tau proteins templates naïve tau monomers that adopt the same misfolded structure to extend the fibril (2–4). Tau species capable of adopting and propagating the tauopathy-specific inclusions are referred to as tau prions or tau strains, while the literature does not precisely define the nature and composition of these species. We henceforth refer to the unique fibril structures with sustainable templated self-propagation capabilities as tau prions (5).

Research shows that tau prions vary in seeding capacity, immunostaining patterns, isoform involvement, cell type specificity, and responses to protease digestion or guanidine denaturation (6–8). Consequently, the specific tau folds associated with each disease may fundamentally influence the course of disease propagation at detailed molecular levels and hence act as highly selective strains even if they differ in seemingly small details in protein folds. The implication is clear: each unique tau structure may require tailored therapeutic and diagnostic strategies that necessitate precision targeting each distinct tauopathy. This discovery underscores the importance of recreating tau prions that adopt the tauopathy-specific folds as targets with high specificity for the development of therapeutics and biomarkers (9, 10). To date, postmortem human tissue-derived tau aggregates are still the gold standard target for tauopathy research, but these samples are scarce and heterogeneous in composition (11, 12). Synthetic reconstruction of tau fibrils from recombinant proteins is a grand challenge, but necessary for generating a sustainable, scalable, and impurity-free alternative target to brain derived tau fibrils (13).

For decades, various approaches have been employed to produce tau fibrils. One commonly used approach to generate tau fibril has been the addition of polyanionic cofactors, with heparin or RNA, being the most prevalent ones. However, structural studies have revealed that full length tau fibrils formed by these cofactors *in vitro*, unless other conditions are imposed, differ significantly from those found in tauopathy disease states (14, 15). A recent cryo-EM study has demonstrated that a carefully selected tau construct, dGAE (tau297-391), can adopt Alzheimer’s Disease (AD) and Chronic Traumatic Encephalopathy (CTE)-like conformations *in vitro* in the presence of different salt and buffer conditions (16). Nonetheless, the underlying mechanisms and kinetics governing the success of these specific conditions remain unclear, leading to the well documented practical challenge that inconsistent results are found across laboratories (12, 17, 18). Another approach is to rely on brain derived fibrils as seeds to generate fibrils from monomeric tau by cellular or *in vitro* seeding; these approaches generated AD-like structures verified by cryo-EM, but reportedly produced low fibril quantities or fibrils lacked seeding competency (19, 20) thereby limiting their utility as pathological tau models. Studies have suggested that pseudo-phosphorylation at specific sites of full-length tau narrow the tau folding pathway towards tauopathy-mimicking fibrils (21), but this has not yet resulted in robust and generalizable solutions that can be built upon, likely because the design principle of selecting specific post translational modification combination is not known. Hence, the reliable replication of full disease cores remains an unresolved challenge which arises from the inherent difficulty in directing a long, intrinsically disordered protein to adopt a specific fold among numerous energetically similar conformations.

Here, we present a novel strategy involving the use of structure-specific mini-prions as model templates to catalyze the formation of critical protein folds *en route* to tauopathy fibrils. This approach is grounded in the hypothesis that fibril seeding is initiated by misfolding of a localized critical motif that acts as a nucleating event to lower the kinetic barrier for the association and folding of the tau protein to form the target fibril core structure. Our design strategy is to identify a small part of the target fold whose formation we hypothesize to be a critical, activation energy lowering, folding event for the target fibrillization pathway, and design a short peptide that folds and stacks to form fibrils that contain the target fold. This peptide that we refer to as a mini-tauopathy motif, due to their limited number of amino acids, explores a significantly reduced conformational landscape that is computationally accessible and can be further manipulated through the choice of sequence, mutations or modifications to adopt a homogeneous target fibril structure. The careful selection of a mini-prion motif that encapsulates the essential properties of a complete fibril core is central to our approach. The most amyloidogenic sequences critical for tau fibrillization are the PHF6 (^306^VQIVYK^311^) and PHF6* (^275^VQIINK^280^) motifs that have been shown to form fibrils with self-complementary steric zippers according to micro-electron diffraction, but without prion-like seeding competency (22–24). Our design approach is distinct in that we link the PHF6 segment to a peptide segment that forms a stabilizing ‘counterstrand’ to ensure that it adopts a fold with its side chains and β-sheet forming hydrogen bonds oriented as found in pathological tau fibrils. We demonstrated this with a peptide encompassing the PHF6 region, paired with a stabilizing ‘counterstrand’ connecting the R2 and R3 junction of tau termed jR2R3-P301L that adopts a U-shape conformation resembling a strand-loop-strand (SLS) motif common to all 4R tauopathy fibrils (25). These peptide fibrils also displayed prion-like competency for self-seeding and seeding larger tau with 4R isoform-specificity *in vitro* and in biosensor cells (25, 26). Nevertheless, the mini-4R tauopathy prions did not populate a uniform target conformation of 4R tauopathies (25) and the seeded tau fibrils did not form high quality homogeneous fibrils amenable for structure confirmation (26).

In this study, we report on the design of a mini-tauopathy sequence to mimic a critical minimal tau folding motif found in mixed 3R-4R tauopathy fibrils of AD and CTE that, when patient-derived, show prion-like seeding competency. For simplicity, we refer to this as the mini-AD peptide and the fibril formed of the mini-AD peptides as mini-AD prions that were designed to contain the PHF6 segment and its counterstrand in AD or CTE fibrils. The significance of the mini-AD motif is that it is not only contained in AD and CTE fibrils, but also in the fibrillar intermediate structures proposed to be *en route* to mature AD or CTE fibrils. Hence, the mini-AD motif offers a common target that encompasses AD, CTE and other fibrillar intermediates along the AD/CTE fibrillization pathway.

We confirm that mini-AD peptides readily form homogeneous fibrils, while a combination of continuous wave Electron Paramagnetic Resonance studies (cw-EPR), cryo-EM and computational predictions suggest that the mini-AD fibrils adopt the targeted disease structure-specific intramolecular motif. These mini-AD fibrils are also competent for seeding tau ten to twenty times its size, *in vitro* and in cellular models. Using a combination of DEER and cryo-EM, we successfully tracked the structural evolution of dGAE fibril formation seeded with mini-AD fibrils under different seeding conditions known to favor AD vs CTE fibrils. This study showcases the potential of disease fold-specific synthetic mini-prion motifs as next generation targets to aid the development of therapeutic and diagnostic strategies, and present design concepts that can be extended to other tauopathy fibril structures whose formation relies on templated aggregation.

## Results

### Design and fibrillization of Mini-AD prion motif

The primary objective of this study was to rationally generate a mini-tau fibril model that adopts a critical structural motif of AD and CTE fibrils and displays prion-like seeding competencies. In AD and CTE, tau fibrils adopt a characteristic intramolecular fold spanning residues 306–378 of the tau sequence (Figure 1a). This region folds into a two-layered structure that adopts either an elongated, open C-shaped conformation in CTE or a more compact, C-shaped conformation in AD (Figure 1b) (27, 28). Additionally, the CTE fold contains a hydrophobic cavity that encloses non-tau densities, speculated to be ions (16). We hypothesized that the steric zipper-like motif formed by the PHF6 sequence (^306^VQIVYK^311^) and its counter strand (^374^HKLTF^378^) that constitute the beginning and end of the tau segment forming the AD and CTE fibril core (Figure 1b) serve as a critical mini-prion motif. To generate this mini prion, a chimera peptide was designed by connecting these sequences by a flexible linker (PGGGN) (Figure 1c). PGGG type sequences are present between each repeat domain in the microtubule binding region (MTBR) (29). The proline residue helps bend the backbone and locally disrupt the secondary structure, which, in combination with the flexible GGG linker favors the formation of U-shaped hairpin motifs seen in several known tauopathy fibril structures. This 16-residue chimera that we refer to as a first generation mini-AD peptide was hypothesized to form a U-shaped intramolecular hairpin motif upon fibrillization, with the expected steric zipper formed by stabilization of PHF6 by the counter strand ^374^HKLTF^378^. This hypothesis was further supported by Alphafold2 (AF2) and Alphafold3 (AF3) simulations predicting the expected mini-AD structure (Figure 1c, Figure S1).

**Figure 1.**
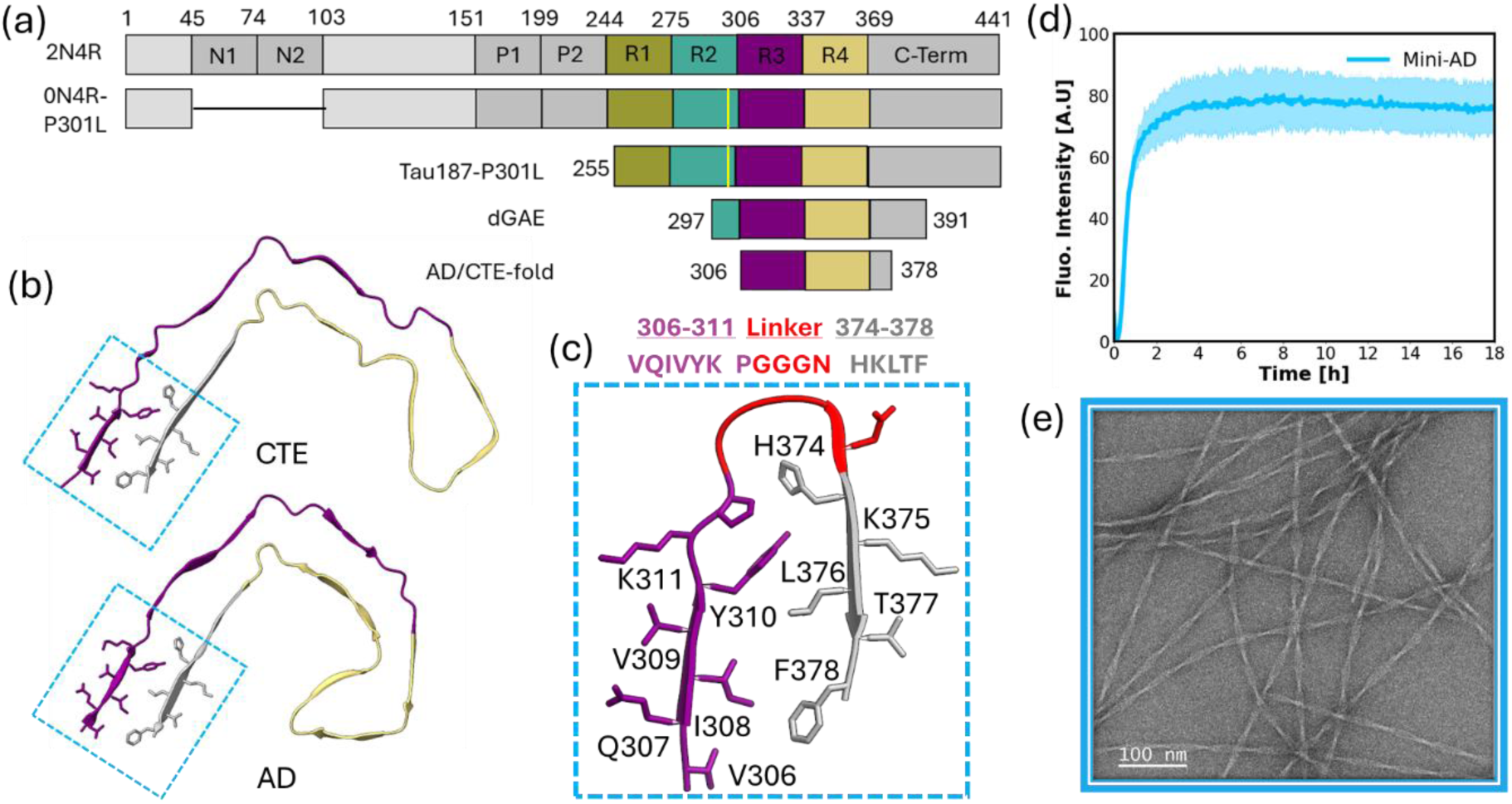
(a) Schematic overview of tau constructs used in this study. The top panel shows the domain organization of the full-length 2N4R isoform of tau with distinct color coding for the N-terminal inserts (N1, N2), proline-rich regions (P1, P2), microtubule-binding repeats (R1–R4), and the C-terminal domain. The constructs used in this study— 0N4R-P301L, Tau187-P301L (with the P301L mutation indicated by a yellow mark), and dGAE—are shown below, aligned with the same domain color scheme. Regions corresponding to the sequences that form Alzheimer’s disease (AD) and chronic traumatic encephalopathy (CTE) fibril folds are also indicated. (b) Cryo-EM-derived intramolecular folding of tau in CTE type I (PDB: 6NWP) and AD paired helical filament (PHF) folds (PDB: 5O3L), with the mini-AD motif highlighted. (c) Amino acid sequence of the mini-AD peptide and its predicted structure from AlphaFold2. (d) Thioflavin T (ThT) fluorescence assay showing the fibrillization kinetics of mini-AD. (e) Negative-stain TEM (ns-TEM) images of mini-AD fibrils, revealing twisted fibrillar morphology (Scale bar: 100 nm).

Next, we tested the fibrillization propensity of the peptide. The mini-AD peptides were fibrillated in the presence of heparin in 20 mM HEPES buffer, at 37 °C and shaking at 200 rpm, and aggregation kinetics were monitored by Thioflavin T (ThT) binding assay. The ThT binding curve showed no lag time, and the reaction plateaued within 4 hours of adding heparin (Figure 1d). The formation of fibrils was further confirmed by negative stain TEM (ns-TEM) showing a homogeneous distribution of long helical filaments (Figure 1e). While adding heparin as a cofactor was essential for readily forming large quantities of high quality mini-AD fibrils, heparin is not expected to alter the expected fold. Instead, heparin stabilizes the long-range stacking of tau fold that this peptide intrinsically favors by providing counter anions that relieve the “charge frustration” of the vertically stacked lysines residues (K311, K374). In fact, negatively charged cofactors such as heparan sulfate proteoglycans (HSPGs) have been proposed as natural cofactors associated with tau aggregation and/or tau prion propagation (30). Our primary goal is to achieve the intra-molecular fold found in AD or CTE fibrils within the mini-AD fibril construct. This target fold, if adopted, is accessible at the fibril end surface and is expected to dictate templated aggregation. We next focus on the structure determination of the peptide fibrils.

### Structural characterization of mini-AD fibrils

We next determined the conformational distribution of the mini-AD fibrils to test whether the expected structural motif is the dominant population. We employed site-directed spin labeling (SDSL) combined with electron paramagnetic resonance (EPR) spectroscopy. A cysteine residue was introduced at each terminus of the mini-AD chimera peptide, followed by spin labeling with MTSL (S-(1-oxyl-2,2,5,5-tetramethyl-2,5-dihydro-1H-pyrrol-3-yl)methyl methanesulfonothioate). For short peptide fibrils, end-to-end spin labels tend to be minimally perturbing for the steric zipper interaction. AF2-predicted fibril structures containing the mini-AD fold suggests an inter-label distance of 0.75 - 2.25 nm, (Figure 2a) a distance range that would give rise to both spin exchange and through-space dipolar interactions leading to EPR spectral broadening (25). Hence, we chose cw-EPR line shape analysis conducive to distance characterization below 2 nm for nitroxide spin labels.

**Figure 2.**
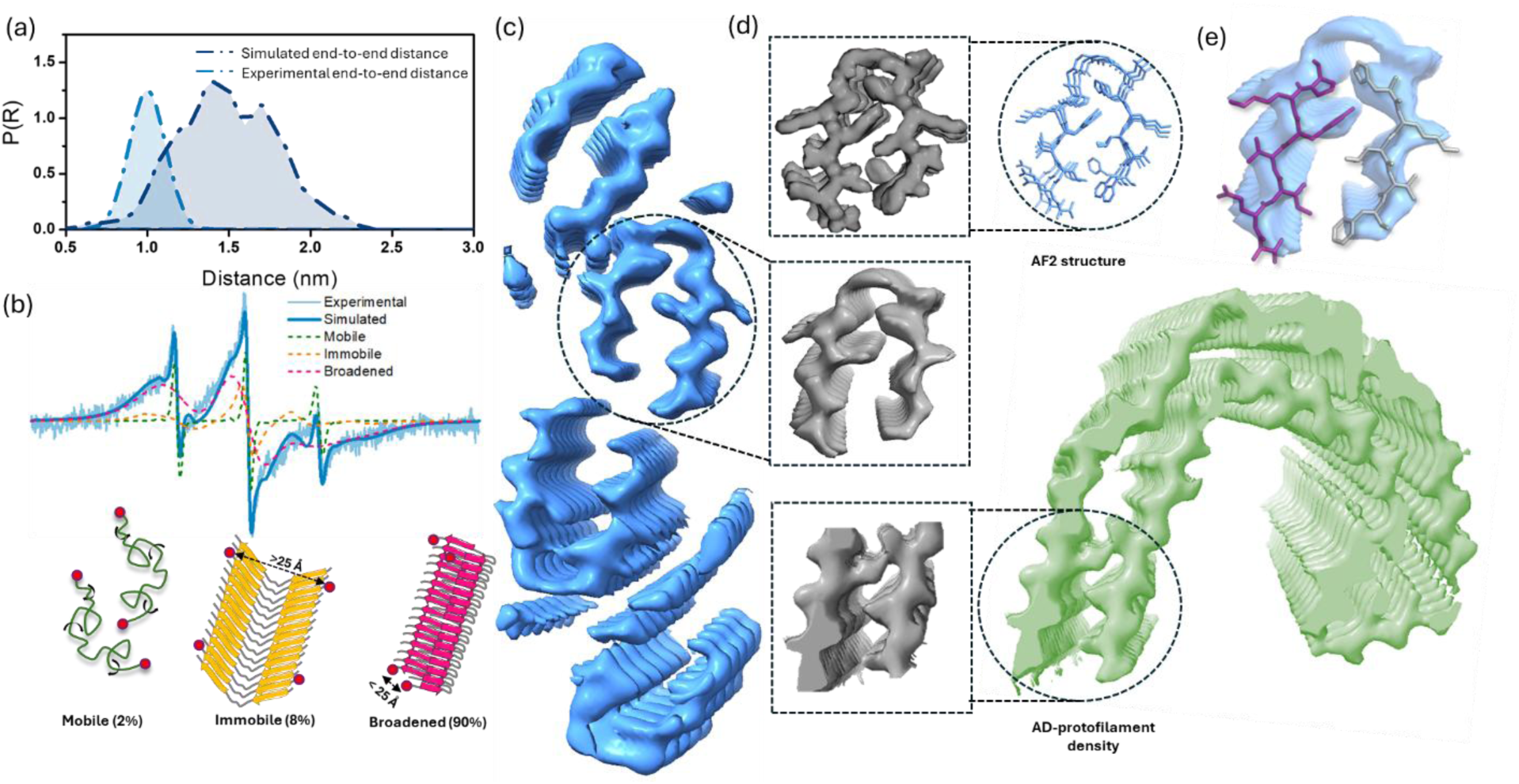
(a) Distance distribution between spin labels at the termini of the mini-AD fold: prediction from the AF2 model using DEERPREdict (dark blue) versus experimental distribution from cw-EPR measurements of spin-labeled mini-AD fibrils analyzed with ShortDistances (cyan). (b) Top: cw-EPR spectrum of mini-AD fibrils showing the experimental signal (pale cyan), simulated fit (dark cyan), and contributions from distinct spectral components: mobile (green, 2%), immobile (yellow, 8%), and broadened (bright-pink, 90%). Bottom: Corresponding structural models for each spectral component are shown using the same color scheme, with red dots indicating spin label positions and annotated end-to-end distances. (c) Cryo-EM density map of mini-AD fibrils, viewed along the fibril axis, revealing a dimer-of-dimers architecture composed of four U-shaped mini-AD folds. (d) Comparison of the cryo-EM density map for a single mini-AD unit (middle) within the cryo-EM density map of mini AD with (i) a simulated 2.4 Å resolution map (top) of the AF2-predicted trimer structure and (ii) the corresponding density (bottom) of an AD-like fibril protofilament assembled *in vitro* (pale green density, PDB: 7QKK, shown as a representative map for all AD/CTE-like fibrils containing mini-AD motif) (e) sidechain orientation of the PHF6 region and ^374^HKLTF^378^ sequence of AF2 structure of mini-AD overlaid on the cryo-EM density of a single mini-AD fold suggesting cross-β interactions as predicted.

To minimize contributions from intermolecular spin–spin interactions, fibrils were assembled by co-incubating 5% doubly labeled peptide with 95% unlabeled peptide. After 24 hours of fibrillization, unincorporated monomers were removed, and cw-EPR spectrum were recorded for the fibrils. Spectral deconvolution using previously established methods (31) revealed three components: (1) a mobile component from freely tumbling spin labels (residual monomers), (2) an immobile component corresponding to non-interacting spin labels incorporated into fibrils, but separated from the next nearest spin label by >2.5 nm, and (3) a dominant broadened component indicative of spins within <2.5 nm—consistent with a U-shaped fold wherein the peptide termini are in close proximity (Figure 2b).

Quantitative analysis showed that ∼90% of the spin labels contributed to the broadened signal, representing spin-exchanged and/or dipolar broadened components, suggesting that the vast majority of the mini-AD peptides adopt the expected fold within fibrils. The mobile and immobile components accounted for ∼2% and ∼8%, respectively. Semi-quantitative analysis of distance distribution corresponding to the broadened component was calculated using the ShortDistances software (32), showing that a major distance population is centered between 0.5 – 1.5 nm (Figure 2a). Minor population of distances exceeding 1.5 nm may still be present but are harder to quantify by cw-EPR line shape analysis since the intrinsic nitroxide linewidth is of the same order of magnitude as the dipolar broadening of spin labels at >1.5 nm separation. The experimentally observed distances align well with the computed distance distribution from the AF2-predicted fibril structure (Figure 2a). These findings confirm that ∼90% of mini-AD fibrils adopt a U-shaped conformation with closely apposed termini.

We next performed cryo-EM to determine the overall structures of the mini-AD fibrils. Two-dimensional (2D) classification analysis showed that a major (70-75%) population form helical fibrils which resembled a paired filament with a crossover distance of 70 nm, as well as minor populations of straight and twisted forms which were not further characterized, indicating minimal heterogeneity in the sample (Figure S2a,b). After multiple rounds of 3D classifications, we resolved a class that is comprised of 4 distinct protofilament subunits, as seen in the cross-sectional view, that each exhibit the predicted shape of the mini-AD hairpin (Figure 2c,d, S2c). These are arranged in a dimer-of-dimers configuration, where two U-shaped mini-AD folds form a dimer that further associates with another such dimer to form a tetrameric assembly (Figure 2c, S2c). Other classes showed a similar overall structure but with poorer resolution (Figure S2c). Further refinements did not converge to a higher-resolution structure, potentially due to heterogeneity in the β-sheet stacking and flexibility of the linker region. Thus, accurate modeling of the mini-AD peptide was not feasible. Nonetheless, the overall cross-sectional view of the protofilaments matches the predicted mini-AD hairpin arrangement (Figure 2d), indicating the cross-β interactions for these residues are likely similar to what is observed in the AD and CTE folds (Figure 2d,e).

### Mini-AD seeds catalyze AD/CTE-like fibrillization pathway aided by salt effects

Next, we proceeded to test the ability of mini-AD seeds to induce fibrillization of larger tau proteins *in vitro*. We selected a tau construct spanning amino acids 297 to 391 (Figure 1a) known as dGAE as the tau substrate to be seeded (16, 33). This construct encompasses the full AD- or CTE fibril core and has been reported to adopt these structures under different fibrillization conditions determined by the choice of salt and buffer (16). However, the resulting fibril structures are highly sensitive to subtle variations in experimental conditions of what appears to be a kinetically driven aggregation reaction (17, 18, 21).

Our own attempt to initiate aggregation of dGAE without seeds, under conditions similar to that described by Lövestam et al.(16), that generates AD-like fibrils, was unsuccessful. We used recombinantly expressed and purified dGAE (See Materials and Methods) with the natural cysteine at site 322 mutated into serine to prevent formation of disulfide linkages (34). No increase in ThT fluorescence was observed when incubating 200 μM dGAE in 10 mM Phosphate Buffer (PB), 10 mM DTT in the presence of 200 mM and 400 mM MgCl_2_ at a shaking speed of 200 RPM at 37°C (corresponding to reaction conditions reported to generate AD-like fibrils (12, 16), referred to as AD-reaction henceforth) even after 60 hours of incubation (Figure S3a). The absence of fibrils was further confirmed by ns-TEM (Figure S3b). When changing the reaction condition to incubating 200 μM dGAE in 50 mM PB and 400 mM NaCl with all the other conditions remaining the same (referred to as CTE-reaction), an increase in ThT signal was observed after 40 hours (Figure S3a). This was further confirmed by ns-TEM showing a heterogeneous population of fibrils (Figure. S3c). These studies confirmed that the CTE reaction more readily proceeds than the AD-reaction. Since 400 mM salt concentration gave some success with fibril formation, we chose to use this salt concentration for the seeding studies for both AD and CTE reactions.

We hypothesized that fibrilization of dGAE to AD-like structures is a kinetically limited reaction that sensitively depends on a myriad of environmental factors that may affect the reaction pathway and hence would require a catalyst to readily proceed. Along similar lines, Lövestam et. al. reported that the formation of a first intermediate amyloid (FIA) structure is the crucial rate limiting step in the selection of the AD-like fibrilization pathway that, once formed, can more readily nucleate the formation of other intermediates (35). Notably, dGAE folds proposed to form *en route* to AD or CTE-like fibrils that come after FIA encompasses the steric-zipper motif across ^306^VQIVYK^311^ and ^374^HKLTF^378^ without any change in its conformation (35). We hence test our hypothesis that mini-AD fibril seeds serve as a catalyst, *in lieu* of FIA formation, to accelerate the aggregation pathways to form AD- or CTE-like fibrils. We propose that the end surface of a mini-AD fibril associates with dGAE monomers along segments encompassing the ^306^VQIVYK^311^ and ^374^HKLTF^378^ region that lowers the kinetic barrier for the formation of fibrillar intermediates containing the mini-AD structural motif (Figure 3a). We believe that the quaternary association of the mini-AD motif has limited influence on templating that is primarily governed by the intramolecular protein fold accessible at the fibril end surface, consistent with evidence from the literature (35, 36). Therefore, discussion on structure propagation will focus primarily on intramolecular folding of both the seeds and the seeded fibrils.

**Figure 3.**
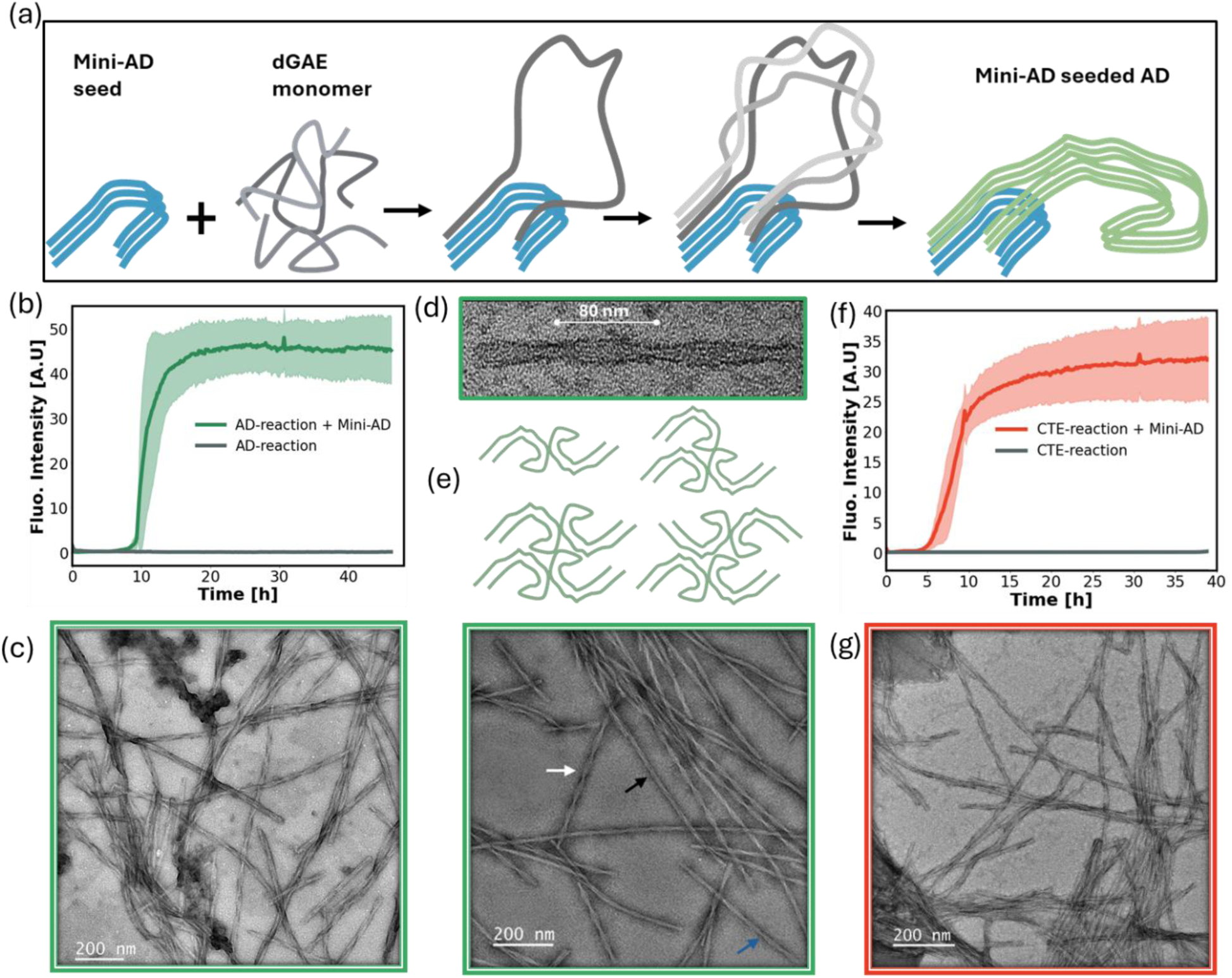
(a) Schematic illustrating the proposed mechanism by which mini-AD seeds promote the templated aggregation of dGAE monomers into AD-like fibrils. Sky-blue denotes stacked arrangement single hairpin structural unit of the mini-AD seed, while pale green represents the full fold adopted by dGAE. (b) ThT fluorescence kinetics for the AD-reaction in the absence (grey) and presence (green) of mini-AD seeds. (c) Representative ns-TEM image of fibrils formed in the AD-reaction seeded with mini-AD seeds (AD-dGAE_ms_). (d) Paired helical filament (PHF) morphology of AD-fibrils, characterized by an ∼80 nm crossover distance. (e) (Top) Schematic depiction of examples of protofilament packing arrangements adopted by AD-like folds of dGAE fibrils(16). (Bottom) Ns-TEM image of AD-dGAE_ms_ with distinct arrows highlighting the observed morphological heterogeneity: white arrows indicate PHF-like structures, while black and blue arrows denote other higher-order assemblies such as THF- or QHF-like morphologies. (f) ThT fluorescence kinetics for the CTE-reaction with (red-orange) and without (grey) mini-AD seeds. (g) Corresponding ns-TEM image of mini-AD–seeded fibrils formed under CTE reaction conditions (CTE-dGAE_ms_). 200 nm scale bars are shown in each ns-TEM image.

To minimize any surface catalyzed secondary nucleation events and to maximize the interaction of monomers with the fibril ends, the long fibrils were broken down into smaller fragments (50-100 nm long) by sonication (Figure S4). Since the mini-AD seed motif is common to both AD and CTE folds (Figure 1b), we hypothesized that the reported salt effects still influence the pathways favoring AD vs CTE folds. In the presence of 50% (molar ratio) mini-AD seeds, the AD-reaction exhibited a ThT fluorescence increase at approximately 8–10 hours, whereas no aggregation was observed in the unseeded reaction over 60 hours (Figure 3b, S3a). In the CTE reaction, the addition of mini-AD seeds triggered fibrillization around 5 hours, nearly seven times faster than the ∼40-hour lag time observed without seeding (Figure 3f, S3a). Critically, the presence of seeds also rendered the reaction robust in terms of reproducibility of the reaction kinetics measured by the ThT time course.

The hypothesis of templated seeding relies on two key assumptions. First, a small fraction of tau filaments, acting as seeds, significantly accelerates fibril formation beyond what would occur naturally without them. Second, the structure of the resulting tau filaments mirrors that of the original seed (2). Since the first criterion is met, we proceeded to evaluating the structure of seeded fibrils.

### Mini-AD seeds different fibril structures *en route* to AD/CTE folds

First, we analyzed the morphological features of the seeded fibrils. AD brain derived fibrils show a paired helical filament (PHF) morphology, with a crossover distance of around 80 nm (Figure 3d). The fibril morphologies of the mini-AD seeded dGAE fibrils in AD reaction (referred to as AD-dGAE_ms_ fibrils henceforth) exhibited heterogeneous fibrils with at least 3 types of morphologies according to ns-TEM, resembling different quaternary association of fibril cores as identified for dGAE fibrils before (Figure 3c, e) including a population of PHF-looking fibrils. It is a known observation as dGAE fibrils tend to form heterogeneous morphologies including triple helical filaments (THFs) and quadruple helical filaments (QHFs) formed by the association of three and four AD-like protofilaments respectively in addition to PHFs (16). It is expected for the AD-dGAE_ms_ fibrils to exhibit heterogeneities as seen with dGAE fibrils, given that the mini-seeds template the VQIVYK-HKLTF fold, hence promoting both AD- and CTE-pathways. The mechanism that controls the quaternary association of these folds should primarily depend on the nature of salt interactions (16) and/or the presence of the fibril’s fuzzy coat (21), not the shape of the mini-prion structure that can only dictate templated extension along the fibril axis. The mini-AD seeded dGAE in the CTE reaction (CTE-dGAE_ms_ fibrils) also generated heterogeneous fibrils, including a major population of a single morphology consisting of thick helical fibrils (Figure 3g).

Despite the heterogeneity in the fibrils, the mini-AD seeded aggregation shows characteristically distinct kinetics and fibril morphologies according to the AD- vs CTE-reaction, suggesting that different structures might be populated. However, the heterogeneous morphologies of the resulting fibrils do not lend themselves to rapid, routine, structure determination by cryo-EM for fibrils generated under a series of different aggregation conditions. To test shape selection in seeded aggregation of dGAE, we turn to Double Electron-Electron Resonance (DEER). Three primary core structures have been reported from cryo-EM studies of dGAE fibrils to form along the AD fibrillation pathway that all contain the mini-AD motif: a C-shaped AD-like fold, an open C-shaped CTE-like fold, and an intermediate J-shaped core structure (IM_J_) (35) (Collectively referred to as AD/CTE-like fibrils henceforth, Figure 4a). Fibrils with these intramolecular folds coexist, likely with folds that are intermediates between these folds, and may associate in various combinations and give rise to diverse quaternary structures and morphological variations across replicates (35). DEER relies on installing a pair of MTSL spin labels on tau monomers to obtain the distribution of intramolecular distances (1.5–8 nm) within a single tau molecule that are sparsely distributed in otherwise non-spin labeled fibrils. We strategically selected residues Q351 and T373 of dGAE for spin labeling, as the intramolecular distance between these sites can effectively distinguish between all three structural variants along the AD/CTE-pathway (Figure 4a).

**Figure 4.**
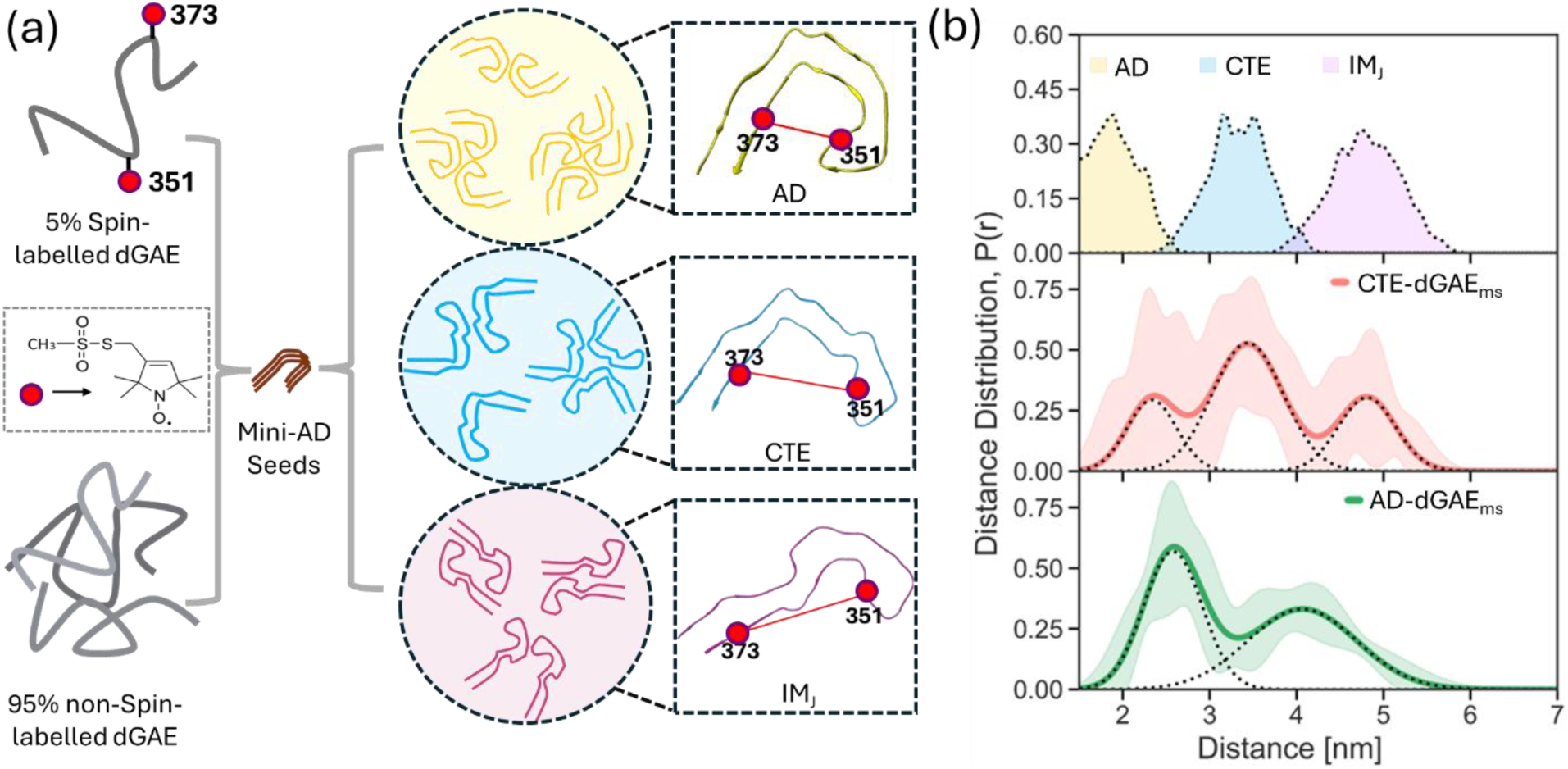
(a) Schematic representation of the DEER experiment used to track structural ensembles seeded by mini-AD. A mixture of 5% spin-labeled dGAE monomers (MTSL labels shown as red dots at positions Q351 and T373) and 95% unlabeled dGAE is incubated with mini-AD seeds. Three major structural classes that can be templated by mini-AD—AD-like (yellow), CTE-like (cyan), and IM_J_-like (pink)—are illustrated, each showing distinct quaternary arrangements and intramolecular folding patterns, with the relative separation between spin-label sites indicated. (b) Simulated inter-spin distance distributions for the three expected structures (top panel: AD-like in pale yellow, CTE-like in pale cyan, IM_J_-like in pale pink) compared to experimental DEER data from mini-AD– seeded dGAE fibrils under CTE-reaction (CTE-dGAE_ms_, middle panel, red-orange) and AD-reaction conditions (AD-dGAE_ms_, bottom panel, green) showing shifts in populations.

A small fraction of the dGAE monomers (5%) were doubly spin labeled and mixed into unlabeled dGAE (95%), such that the distribution of distances measured by DEER captures the population of different intramolecular protein folds. DEER quantitatively reports on the distribution of intramolecular distances, P(r), across all fibril populations encompassing highly heterogeneous quaternary associations. The anticipated P(r) distribution was simulated based on the reported PDB structures of AD, CTE, and IM_J_ fibrils using the DEERPREdict software (37), showing that the three populations are well separated within the DEER measurable distance range (Figure 4b, top).

The P(r) distribution for CTE-dGAE_ms_ showed a broad range with peaks corresponding to all three expected structures (Figure 4b, middle). This suggests that the mini-AD seeded dGAE under CTE conditions forms all three expected structure types, consistent with what was reported for dGAE fibrils formed in the absence of seeding (35). Interestingly, the distribution indicates that a major population adopts a CTE-like fold with minor populations of AD and IM_J_ fibrils (Figure 4b, middle). Remarkably, the distance population of AD-dGAE_ms_ fibrils are shifted towards AD-like folds, with a significant CTE-like population and the IM_J_ fold still present (Figure 4b, bottom).

Although DEER provides insights into the broad distribution of fibril populations, higher-resolution data is needed to clearly determine the presence of the mini-AD motif in the seeded fibrils. To this end, we performed cryo-EM on dGAE tau filaments seeded with the mini-AD fibrils in AD-reaction as described above. 2D classification showed a mixture of filament morphologies, with the majority (∼63%) showing a conformation which resembles the Type I tau filament from patients with CTE (28) (Figure S5a,b), suggesting the population of CTE-like structures. However, further processing is necessary to determine unambiguously whether these filaments are successfully recapitulating the Type I CTE fold.

Together these observations suggests that mini-AD fibrils can seed longer tau proteins encompassing the entire AD/CTE core-forming region to form AD- and CTE like conformations and that the so seeded dGAE fibrils exhibit salt-mediated structural selection similar to salt-induced dGAE fibrils reported before (16, 35).

### Multiple generational seeding using AD/CTE-dGAE_ms_ seeds suggest templated propagation governed by seed structure

While mini-AD successfully catalyzed the formation of AD/CTE-like fibrils and fibril intermediates, the resulting fibrils are a mixture of expected structures encompassing the mini-AD motif, with their relative population influenced by solution conditions. This is expected since mini-AD lacks the complete structural framework of a full fibril seed and hence exerts limited control over the conformation adopted by the rest of the fibril. In contrast, mini-AD–seeded dGAE fibrils (AD-dGAE_ms_) possess a complete fibril core and hence offers a more efficient template to promote structural propagation.

We hypothesized that mini-AD seeding captures early, motif-specific templating events representative of initial disease propagation, involving intermediate structural transitions, influenced by environmental factors like salt effects. On the other hand, full-length seeds like AD-dGAE_ms_ or CTE-dGAE_ms_ recapitulate later stages of fibrillization where specific full folds are amplified. To test this concept, we seeded dGAE monomers with a AD-dGAE_ms_ fibril batch showing significant paired helical filament (PHF) populations under AD-reactions (Figure S6a).

The addition of AD-dGAE_ms_ seeds immediately initiated aggregation that progressed steadily over 24 hours according to ThT fluorescence (Figure 5a). This is in contrast to the seeding kinetics with mini-AD fibrils under AD-reactions that exhibited a distinct lag phase (Figure 3b). The lag likely reflects on the time required for dGAE monomers to interact with the mini-motif and for ThT-active intermediates to emerge. Previous studies have also reported on ThT-inactive intermediate species (35) along the pathway, which could explain this delay. The absence of a lag in AD-dGAE_ms_ seeding suggests that these seeds already possess a more complete structure capable of templated seeding ThT-active fibrils.

**Figure 5.**
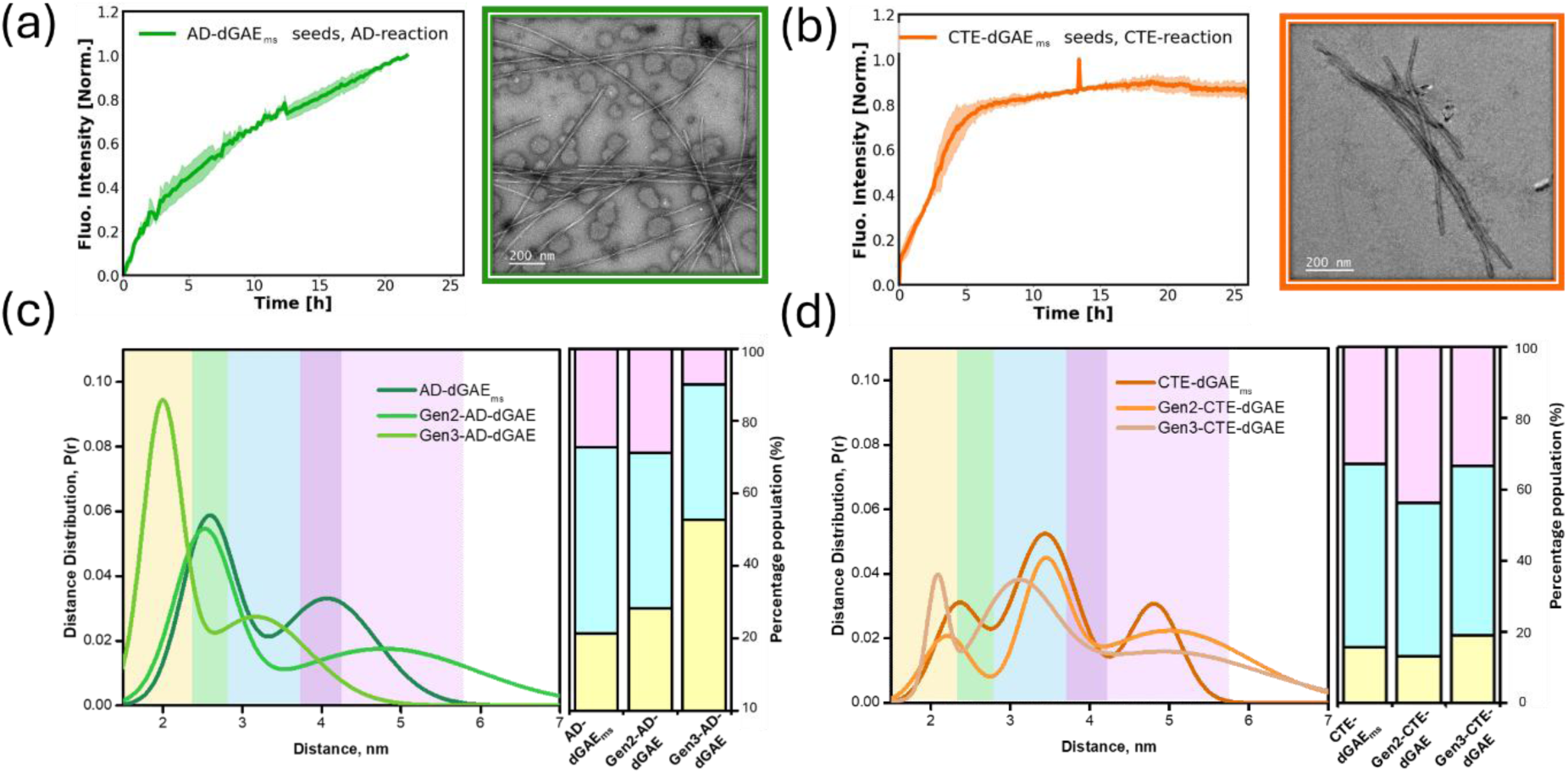
(a, b) Normalized ThT fluorescence kinetics (left) and ns-TEM images (right) of fibrils formed by (a) AD-dGAE_ms_ seeding dGAE monomers under AD-reaction conditions (light green), and (b) CTE-dGAE_ms_ seeding dGAE monomers under CTE-reaction conditions (orange). 200 nm scale bars are shown in each ns-TEM image. (c, d) DEER-derived distance distributions (P(r)) for fibrils generated by multiple rounds of seeding under (c) AD and (d) CTE-reaction conditions. The legend for each P(r) distribution indicates the fibrils for which the distribution is shown. Only the mean distributions are shown; for full datasets including error estimates, see Figures S7-10. Approximate population percentages (bar graphs on the right of the P(R) distribution) were estimated from the relative peak areas within the predicted distance ranges for the three expected structures: AD-like (pale yellow), CTE-like (pale cyan), and IM_J_-like (pale pink). The expected distance distribution range for each structure is shown in the P(r) distribution plots with the same colors. The overlapping regions of expected distances for AD and CTE is shown in pale green and that of CTE and IM_J_ are shown is dark pink in the P(r) distribution plots.

A gradual, steady increase of β-sheets over 24 hours is indicative of a slow elongation process consistent with monomer addition at fibril ends, characteristic also shown by the PAD12 tau seeded with brain-derived AD fibrils (21) reported recently. Fibrils collected after 24 hours (Gen2-AD-dGAE) showed enhanced structural homogeneity with further enriched PHF content, a result confirmed in triplicates (Figure 5a, S6a). Further propagation (Gen3-AD-dGAE) maintained this trend, yielding consistent fibril morphology and increased PHF content (Figure S6a).

DEER analysis revealed a progressive enrichment of AD-like folds across successive seeding generations, accompanied by a corresponding decline in CTE- and IM_J_-shaped conformers (Figure 5c). The distance distributions obtained from dGAE fibrils seeded over multiple generations indicate that AD-dGAE_ms_ consist of two distinct structural populations corresponding to the AD-like folds: one centered at ∼2.6 nm, which falls between the expected distances for AD and CTE folds, and another at ∼2.0 nm, which becomes enriched and dominant only in Gen3-AD-dGAE fibrils (Figure 5c). This suggests the presence of intermediate conformations during early seeding time points, potentially adopting structural features between the compact C-shaped AD fold and the open C-shaped CTE fold. The cryo-EM structures of dGAE fibrils formed along the AD/CTE-like pathway support this concept as their structures reveal subtle conformational shifts along a structural continuum from compact to open folds (35). The canonical AD and CTE folds exhibit mean distances of ∼2.0 nm and ∼3.4 nm, respectively, across spin-labeled sites Q351C and T373C, and hence we suggest that the ∼2.6 nm peak represents a partially open AD-like conformation (Figure 5c). These results support a mechanism wherein specific folds are selected and amplified over successive seeding rounds, resembling the *in vivo* progression toward a dominant fibril conformation, as observed in postmortem brain-derived samples.

A parallel experiment using CTE-dGAE_ms_ fibrils showed that seeding under CTE conditions resulted in a dramatically different kinetic profile with faster ThT signal increase that plateaued within six hours (Figure 5b). The resulting fibrils closely resembled the original CTE-dGAE_ms_ morphology according to ns-TEM (Figure 5b, S6b). Notably, DEER analysis of Gen2 and Gen3-CTE-dGAE fibrils exhibited P(r) profiles consistent with the CTE structural motif and do not evolve much (Figure 5d). This suggests that the CTE fold forms more readily and CTE-dGAE_ms_ faithfully templates dGAE monomers.

The contrasting kinetics and structural outcomes observed upon seeding dGAE with AD-dGAE_ms_ and CTE-dGAE_ms_ prompted us to explore the role of environmental conditions in templating fidelity. We investigated whether altering the salt composition would affect propagation in templated aggregation. Seeding dGAE monomer under CTE-reaction conditions using AD-dGAE_ms_ fibrils produced a ThT kinetic curve and fibril morphology resembling the results obtained when the same seeds were used to seed under AD-reaction conditions (Figure S11a). The DEER derived P(r) distribution also matches that of AD-dGAE_ms_ fibrils, indicating that the templating action of the seed dominates the structural outcome (Figure S11c).

Seeding dGAE monomers under AD-reaction conditions using CTE-dGAE_ms_ fibrils produced, again, a ThT kinetic curve and fibril morphology governed by the nature of the seeds irrespective the salt used (Figure S11b). The P(r) of DEER showed a major peak around the expected distance for CTE (Figure S11c), consistent with the seeds used. In both cases, minor populations corresponding to alternate folds remain, showing that there still is some environmental influence on the seeded propagation.

These results establish that mini-AD seeding generates a range of representative structures along the AD/CTE pathway that are themselves seeding competent templates. Once a complete stable seed structure is formed, it dominates the conformation selection of the templated fibrils, bypassing intermediate and environment-sensitive transitions. Thus, mini-AD seeding more accurately models early-stage pathway selection, while seeding with AD-dGAE_ms_ and CTE-dGAE_ms_ offer robust models for later-stage templated propagation.

### Mini-AD seeds induce mixed 3R-4R tau fibrils in cells like AD-patient brain fibrils

To evaluate the seeding capability of mini-AD in a cellular context, we utilized H4 cells stably expressing mClover3-tagged 0N4R-P301L tau. The P301L mutation is known to promote tau aggregation and has been previously shown to be required for puncta formation upon exposure to exogenous fibrils (25, 26). Treatment of the cells with mini-AD seeds resulted in robust puncta formation (Figure 6a), demonstrating that mini-AD can effectively seed aggregation of the 0N4R tau isoform—an endogenous human brain isoform that is approximately twenty times larger than the mini-AD construct.

**Figure 6.**
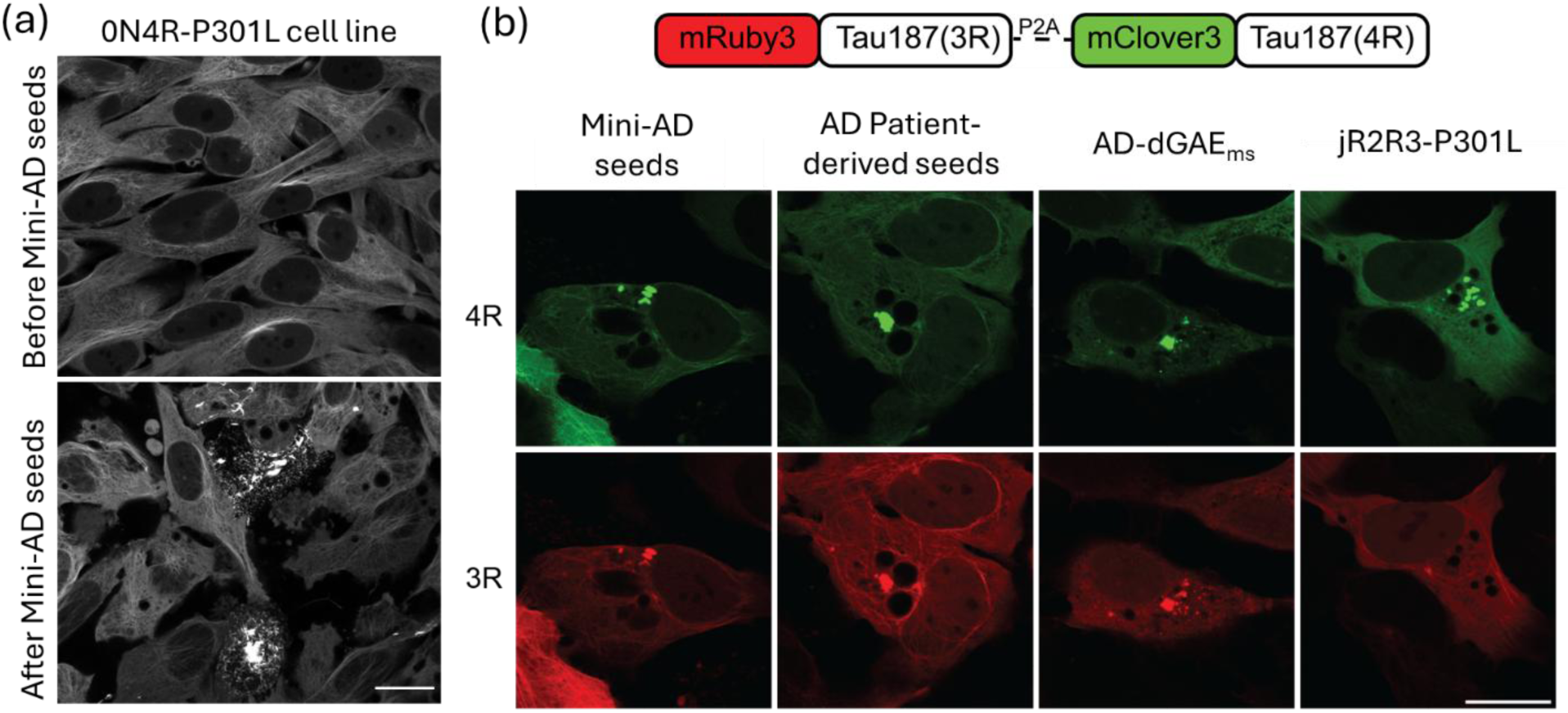
Tau aggregate formation, propagation, and their characteristics in cells seeded with mini-AD seeds. (a) H4 cell stably expressing mClover3-0N4R-P301L, seeded with mini-AD fibrils, imaged at 0- and 12-h post addition of mini-AD seeds (Scale bar 20 μm). Puncta formation is observed in the latter. (b) (Top) Schematic representation of the plasmid construct used to express two copies of Tau187. The construct includes mRuby3-Tau187(3R) (red), followed by a self-cleaving P2A sequence, which enables expression of mClover3-Tau187(4R)-P301L (green) within the same cell. (Bottom) Confocal fluorescence images of H4 neuroglioma cells following fibril treatment. Cells expressing the Tau187 dual-isoform construct were incubated for 24 hours with mini-AD fibrils, patient-derived AD fibrils, AD-dGAE_ms_ or jR2R3-P301L fibrils. Mini-AD, AD-dGAE_ms_ and patient-derived AD fibrils induced puncta formation in both the 3R (green) and 4R (red) channels, demonstrating incorporation of both tau isoforms. In contrast, jR2R3-P301L fibrils selectively seeded the 4R isoform, with no detectable puncta formation in the 3R channel (Scale bar: 20 μm).

To assess the isoform seeding specificity of mini-AD fibrils, we utilized our previously described mammalian cell model expressing two Tau187 isoforms: mClover3-Tau187(3R) (3-repeat isoform) and mRuby3-Tau187(4R)-P301L (4-repeat isoform) (26). Following 24 hours of mini-AD fibril treatment, puncta formation was observed in both the 3R and 4R Tau187 fluorescence channels (Figure 6b). Notably, these signals were always colocalized, indicating that both 3R and 4R Tau187 isoforms were incorporated into the same fibril-seeded aggregates.

As a positive control, we tested patient-derived Alzheimer’s disease (AD) fibrils, which are expected to seed both 3R and 4R tau. As anticipated, robust puncta formation was observed in both channels (Figure 6b), further validating that patient-derived fibrils promote aggregation of both tau isoforms. Furthermore, we tested the isoform selective seeding activity of AD-dGAE_ms_ seeds, which showed very similar trends (Figure 6b).

Finally, as a negative control known to seed only 4R isoforms, we introduced jR2R3-P301L fibrils, which are known to mimic 4R tauopathies (25, 26). In contrast to the mini-AD and patient-derived fibrils, puncta formation was restricted to the 4R Tau187 (mRuby3) channel, with no detectable aggregation in the 3R Tau187 (mClover3) channel (Figure 6b), consistent with 4R isoform-specific seeding.

## Discussion

The generation of tau fibrils that mimic disease folds and replicate seeding competency is a major focus of tauopathy research. However, rational tuning and design, as well as the reliable generation of such fibrils with high homogeneity is a grand challenge (13). In this context, the design of shape-specific mini-prions, using peptide sequences derived from the microtubule binding repeat domain (MTBR) of tau, is a new frontier that this study is exploring. The ability of a peptide to serve as a viable mini-prion model depends on two critical criteria. First, the mini-prion must adopt a fibril structure that replicates the amino acid arrangement of a critical motif found in a disease-relevant fold. Second, it must be capable of templating and amplifying the same motif onto naïve tau monomers to generate seeding-competent fibrils, mirroring the prion-like behavior of disease-associated tau fibrils. The first criterion is essential for achieving the second, given that even minor alterations in amino acid interactions within the prion motif can lead to the propagation of unintended fibril structures. Our own previous study on the jR2R3-P301L peptide came close to meeting these requirements, successfully forming an SLS motif characteristic of 4R tauopathies(25) and even demonstrating 4R isoform-selective seeding (26). One of the fibril folds formed by this peptide most closely resembled the limbic-predominant neuronal inclusion body 4R tauopathy (LNT) fold in terms of amino acid orientation within the core, although it didn’t exactly replicate the amino-acid interactions specific to a 4R tauopathy structure.

In this study we presented a mini-prion fibril that mimics a tauopathy-specific folding motif found in fibrils along the AD/CTE-like aggregation pathway. One of the novel design strategy presented in this study is to generate a shape-specific mini-prion by artificially connecting two sequences known to form thermodynamically stable steric zippers in the disease fold through a linker (PGGG), thereby forcing them to adopt the target mini motif. We demonstrate that the designed mini-AD prion catalyzes the formation of AD/CTE fibrillization pathway intermediates through templated seeding. This finding supports the stepwise templating mechanism proposed for the AD fibrillization pathway (35), in which aggregation begins from a key motif that drives the selection of AD/CTE-like structures. DEER structural studies further reveal that shape propagation is dictated by the core structure of the seed itself, once a final seed core is established, while environmental factors have a smaller influence on its propagation. Our results establish mini-AD as a crucial motif in determining pathway selection for AD/CTE-like fibril formation and introduce mini-AD fibrils as a valuable alternative model for studying AD/CTE-like fibrils.

The development of structure-specific binders that target the unique motifs in the fibrils to inhibit or detect tau aggregation is a major focus in tauopathy research. These binders— including small-molecule binders (38), nanobodies (39–44), antibodies (45, 46), peptide-based binders (23, 47), and positron emission tomography (PET) ligands (48, 49)—are often designed based on screening assays using tau fibrils that do not adopt a disease fold or are devoid of prion-like seeding competency. The screening for small molecule binders often starts with derivatives of existing small molecule binders with non-ideal specificity or that are based on imperfect models for the target structure and surface property, leading to suboptimal binding specificity and affinity. For example, the crystal structures of PHF6 and PHF6* revealed self-complementary steric zipper motifs that guided inhibitor design (23), but the inability of these steric zippers to seed fibril growth limits them as model fibrils to discover inhibitors for prion-like propagation.

A fibril serving as a mini-prion, such as mini-AD, that reliably forms a disease-relevant motif and catalyzes disease pathway-specific seeding offers a platform for designing and testing binders and inhibitors for prion-like propagation, including conformation-specific antibody and nanobody development. The discovery presented in this study offers both conceptual and practical advantages. On one hand, mini-prion fibrils are easier to generate while maintaining high structural fidelity, enabling the reliable synthesis of targets with well-defined structure, morphology, and biophysical properties. On the other hand, a mini motif that captures a key structural region of the full fold provides a powerful tool for screening and refining computationally selected binders from large libraries. This approach enhances specificity by focusing on the relevant epitope, while reducing the risk of false positives that may arise from off-target binding within the full disease-associated fibril structure.

A recent cryo-EM study by Lövestam, Scheres, Goedert and coworkers (35) identified an intermediate species along the *in vitro* fibrillization pathway of dGAE, proposing a stepwise templating mechanism in which the FIA act as an initial nucleation site for the formation of AD/CTE-like fibrils. This finding offers inspiration of how seeding could be initiated by mini-motif intermediate structures, underscoring the importance of targeting them. However, the FIA fibril has not been isolated, and its stability and prion-like properties remain unverified, limiting its utility as a target for AD pathways. Interestingly, FIA fibrils were found to contain a self-complementary steric zipper motif between two PHF6 regions, similar to the one shown by Eisenberg and coworkers (22, 23), which is however not found in the post-mortem fibril fold. The presence of the FIA motif in disease brains has not yet been confirmed and is likely representative of a transient species that forms during the early stages of disease progression, making it more difficult to detect and target. Our results support this, showing that the final seed structure, rather than intermediates like FIA, governs shape propagation once the final disease core forms, as evidenced by the lack of lag time and relatively salt-insensitive behavior of AD-dGAE_ms_ seeding.

We propose that an effective strategy to develop diagnostic and/or therapeutic agents for tauopathies is to target species that persist throughout all stages of disease progression. The study that reports on FIA notes that the residues 305–316 and 370–380 of dGAE, constituting mini-AD adopt the same conformation in all later intermediates following FIA, suggesting that the formation of the mini-AD structural motif is an important event throughout the AD-fibrillization and propagation pathway that selects AD/CTE-like structures. Our observation that mini-AD fibrils seed a diverse population of AD/CTE-like seeded fibril structures further emphasizes that this motif must be common to fibril intermediates *en route* to AD, and CTE fibrils, and hence a promising target.

The design strategy we employed here to generate the mini prion is a generalizable idea that could potentially be extended to other disease folds to generate similar mini prions of other steric zippers. Interestingly, the PHF6 and PHF6* (^275^VQIINK^280^) motifs form conserved steric zipper interactions across various tauopathy-associated fibril structures, interacting with distinct but structurally complementary sequences (50).These interaction partners that stabilize the zipper interactions are well separated from the hexapeptide motifs within the MTBR. For example, the PHF6* sequence also associates with ^373^THKLTF^378^, forming the beginning and the end of the corticobasal degeneration (CBD) fibril folds, similar to the mini-AD motif found in AD and CTE folds. The PHF6 sequence engaging in conserved interactions with ^337^VEVKSE^342^ in Pick’s disease (PiD), CBD, and argyrophilic grain disease (AGD) fibrils is another example. The recurrence of these structurally conserved motifs—formed by well-separated sequences—across multiple tauopathy fibril structures suggests that their formation is unlikely to be coincidental. It is plausible that these conserved interactions serve as key determinants guiding structure selection in different disease-associated fibrillation pathways. The sequential order of such interactions may determine which aggregation pathway is ultimately favored. For instance, the association of PHF6 with ^373^THKLTF^378^ appears to be a critical step that directs fibril templating toward AD/CTE-pathway intermediates following FIA formation. Instead, if the association of PHF6 with ^337^VEVKSE^342^ happens first, it may drive fibril formation toward CBD- or PiD-pathway structures. Furthermore, the subsequent interaction of PHF6*(absent in 3R tau) with ^373^THKLTF^378^ could selectively stabilize a CBD-like fold in 4R tau. Future experimental studies are needed to validate this hypothesis and elucidate the sequence of molecular events governing tau fibrillation pathways. Our study demonstrated that peptide designs that lead to homogeneous mini-prion fibrils are promising strategies to achieve the selection of specific seeding pathways. A combination of such mini-prions specific to unique folds could prove to be alternate targets that are representative of a specific target disease fold for future therapeutic/diagnostic development and offer insights towards a generalized mechanism of templated formation of different disease-relevant tau folds.

## Materials and methods

### Peptide production

All peptide constructs including the spin labelled ones were synthesized by the Peptide Synthesis Core Facility of the Center for Regenerative Nanomedicine at Northwestern University. Peptide identity and purity were confirmed by mass spectrometry and HPLC (>95%). Peptides were hydrated in 20 mM HEPES buffer pH 7.4 to a concentration of 1 mM and were immediately aliquoted and stored at −80 °C until use.

### dGAE expression and purification

Unless otherwise stated, all *in vitro* studies were performed using the N- and C-terminally truncated dGAE construct (residues 291–397, MW: 10.165 kDa) with a C322S mutation to eliminate the native cysteine. The spin-labeled variant, referred to as Q351C–T373C dGAE, was generated via site-directed mutagenesis by introducing cysteine residues at positions 351 and 373 in addition to the C322S mutation.

We followed a previously reported purification protocol with minor modifications (51). Briefly, the gene encoding the desired dGAE tau construct (residues 291–397) was cloned into a pET-28a vector, incorporating an N-terminal His-tag and a TEV protease cleavage site. The plasmid was transformed into E. coli BL21 (DE3) competent cells (Novagen) for protein expression. Starter cultures were grown overnight at 37 °C in 10 mL Luria Broth (LB) from glycerol stocks or freshly streaked plates. These cultures were used to inoculate 1 L of fresh LB medium, which was incubated at 37 °C with shaking at 200 rpm. When the OD₆₀₀ reached 0.6–0.8, protein expression was induced with 1 mM isopropyl-β-D-thiogalactopyranoside (IPTG, Sigma-Aldrich) for 2–3 hours at 37 °C. Cells were harvested by centrifugation at 10000 rpm for 10 minutes (Beckman J-10) and stored at –20 °C until use. For lysis, frozen cell pellets were resuspended in lysis buffer [50 mM Tris-HCl (pH 7.4), 100 mM NaCl, 0.5 mM DTT, 0.1 mM EDTA, 1 mM PMSF, and one Pierce™ Protease Inhibitor tablet (Thermo Fisher Scientific) per 50 mL]. Lysozyme (2 mg/mL), DNase I (20 µg/mL), and MgCl₂ (10 mM) were added, and the suspension was incubated on ice for 30 minutes. The lysate was then heat-treated at 65 °C for 13 minutes to denature bacterial proteins, cooled on ice for 20 minutes, and centrifuged to remove precipitates. The resulting supernatant was applied to a Ni-NTA affinity column pre-equilibrated with Buffer A [20 mM sodium phosphate (pH 7.0), 500 mM NaCl, 10 mM imidazole, 100 µM EDTA]. After washing with 20 mL Buffer A and 15 mL Buffer B [20 mM sodium phosphate (pH 7.0), 1 M NaCl, 20 mM imidazole, 0.5 mM DTT, 100 µM EDTA], the bound protein was eluted using Buffer C [20 mM sodium phosphate (pH 7.0), 0.5 mM DTT, 100 mM NaCl] with a gradient of imidazole (100–300 mM). Eluted fractions were pooled, concentrated using Amicon Ultra-15 centrifugal filters (MWCO 3 kDa; Millipore Sigma), and desalted into 10 mM phosphate buffer (pH 7.4) using PD-10 desalting columns (GE Healthcare). Protein concentration was determined by UV absorbance at 274 nm using an extinction coefficient of 1400 M⁻¹cm⁻¹ (based on tyrosine content). To cleave the His-tag, the sample was incubated with TEV protease (Sigma-Aldrich, Product number 4455, 5-fold molar excess) and 10 mM DTT at 4 °C for 15–16 hours. The mixture was then passed through Ni-NTA resin to separate the cleaved, tagless dGAE in the flow-through and further eluted with buffer A. The resulting protein was re-concentrated and desalted again into 10 mM phosphate buffer (pH 7.4). For further purification, size-exclusion chromatography (SEC) was performed on a HiLoad 16/600 Superdex 200 pg column (GE Healthcare Life Sciences) connected to a Bio-Rad NGC Quest 10 FPLC system. The column was washed with 130 mL degassed Milli-Q water and equilibrated with 160 mL of degassed working buffer [10 mM phosphate buffer (pH 7.4)] at a flow rate of 0.8 mL/min. The sample (2–5 mL) was loaded, and elution was carried out with 150 mL of the same buffer at 0.6 mL/min. Elution peaks were monitored and compared against monomeric dGAE standards. Fractions corresponding to monomeric dGAE were pooled, concentrated (Amicon Ultra-4, 3 kDa MWCO), and quantified by UV absorbance at 274 nm as described above.

### Mini-AD fibrillization

Mini-AD aggregation assays were performed in 20 mM HEPES buffer (pH 7.4). Reactions were set up in Corning™ 384-well solid black polystyrene microplates (Thermo Fisher Scientific), with each well containing 100 μM mini-AD and 20 μM Thioflavin T (ThT). Aggregation was initiated by the addition of heparin (16 kDa average MW) at a 4:1 molar ratio (mini-AD:heparin). ThT fluorescence was monitored using a Tecan Spark microplate reader (excitation: 440/30 nm; emission: 485/20 nm; number of flashes: 16) at 37 °C over 24 hours with 4.5 mm amplitude, linear shaking. Readings were taken every 3 minutes, with the instrument gain set to 50. Each condition was measured in triplicate, and results are presented as the mean ± standard deviation.

For mini-AD seed preparation, aggregation reactions were scaled up to 5 mL under identical buffer and molar ratio conditions, excluding ThT. A 10 μL aliquot was withdrawn from the reaction right after the addition of heparin, mixed with ThT, and monitored by plate reader to confirm aggregation plateau. Upon completion, fibrils were collected and concentrated using Amicon Ultra centrifugal filters (MWCO 100 kDa; Millipore Sigma), followed by three rounds of buffer exchange with 10 mM phosphate buffer (pH 7.4) to remove residual monomers and heparin. The final sample was concentrated to 500 μL.

A small aliquot was characterized by negative-stain transmission electron microscopy (ns-TEM) to verify that the fibrils were well-dispersed and not clumped. Fibrils were then aliquoted (20 μL) and sonicated for 15–30 minutes (30 kHz, 70% power) in a bath sonicator (Cole-Parmer Elmasonic P30H). Ns-TEM imaging was used to confirm fragmentation of fibrils to an average length of ∼50–100 nm. Once this was achieved, seed aliquots were flash-frozen and stored at –80 °C for future use. The final seed concentration was estimated to be ∼500 μM, assuming near-complete conversion of monomers to fibrils during aggregation.

### dGAE fibrillization with and without mini-AD seeds

Aggregation reactions were conducted in Corning™ 384-well solid black polystyrene microplates (Thermo Fisher Scientific). For each well, the following components were added: phosphate buffer (50 mM for CTE-reaction or 10 mM for AD-reaction, pH 7.4), 20 μM Thioflavin T (ThT), 10 mM dithiothreitol (DTT), 200 μM dGAE protein, and the appropriate salt (400 mM NaCl for CTE-reaction conditions, 200 mM or 400 mM MgCl₂ for AD-reaction conditions, from 5M salt stocks in ultrapure water). For mini-AD seeding reactions, sonicated mini-AD seeds were added at a 1:2 molar ratio (mini-AD monomer equivalents in the seeds: dGAE monomers), with the mini-AD seed concentration estimated based on the number of monomer units within the fibril seeds, as described above. ThT fluorescence was recorded using a Tecan Spark microplate reader (excitation: 440/30 nm, emission: 485/20 nm, number of flashes: 16) over a 60-hour period at 37 °C with 4.5 mm amplitude linear shaking. Data points were collected every 10 minutes. Each condition was measured in triplicate, and fluorescence plots report the mean ± standard deviation. For reactions that were intended for downstream applications (e.g., further seeding) and did not require ThT, a parallel reaction was prepared under identical conditions without ThT. After the completion of the aggregation reaction, the fibril products were imaged using ns-TEM to confirm fibril morphology, and the samples were stored at –80 °C for future use. For multiple generational seeding reactions (i.e., dGAE seeding dGAE), the mini-AD-seeded dGAE fibrils were thawed and sonicated for 5 minutes using the same procedure as described for mini-AD seed preparation. The resulting seeds, referred to as AD-dGAE_ms_ and CTE-dGAE_ms_, were assumed to have the same concentration as the starting monomer (200 μM). These were added at 10% (molar ratio) to fresh dGAE aggregation reactions under identical buffer and salt conditions, using 100 μM dGAE monomer. The aggregation was monitored in Tecan Spark plate reader as described above for 24 hours.

### Negative Stain TEM

Negative-stain TEM grids (200-mesh Formvar-coated copper) were glow-discharged for 45 seconds using a PELCO easiGlow system and used within one hour of treatment. A 4 µL aliquot of the sample was applied to the grid, incubated for 1 minute, and then blotted using Whatman filter paper. The grid was stained by briefly applying 4 µL of 1% (w/v) uranyl acetate, followed by immediate blotting. A second 4 µL aliquot of the same stain was then applied, incubated for 1 minute, and blotted again before air drying. Imaging was performed on a JEOL 1400 Flash transmission electron microscope operated at 120 kV accelerating voltage.

### Cryo-EM Sample Preparation and Data Collection

#### Mini-AD

Filaments (3 uL) at a concentration of 200 μM, after fibrillization reaction plateaued, were added to 400 mesh 1.2/1.3R Cu Quantifoil grids which had been glow discharged for 40s. After 15 seconds, grids were blotted for 4 s at 22 °C and 95% humidity using a FEI Vitrobot Mark IV, followed by plunge freezing in liquid ethane. Super-resolution movies were collected at a nominal magnification of 105,000x (physical pixel size: 0.417 Å/pixel) on a Titan Krios (Thermo Fisher Scientific) operated at 300 kV and equipped with a K3 direct electron detector and BioQuantum energy filter (Gatan, Inc.) set to a slit width of 20 eV. A defocus range of −0.8 to −1.8 um was used, with a total exposure time of 2.024 seconds fractionated into 80 0.025-second subframes. The total dose for each movie was 46 electrons/Å^2^. Movies were motion corrected using MotionCor2(52) in Scipion (53) and were Fourier cropped by a factor of 2 to a final pixel size of 0.834 Å/pixel. Motion corrected and dose-weighted micrographs were imported into RELION 5 (54) for further image processing.

#### dGAE seeded with mini-AD

Filaments (3 uL) at a concentration of 200 μM, after fibrillization reaction plateaued, were added to 400 mesh 1.2/1.3R Cu Quantifoil grids which had been glow discharged for 40s. After 30 seconds, grids were blotted for 5 s at 22 C and 95% humidity using a FEI Vitrobot Mark IV, followed by plunge freezing in liquid ethane. Super-resolution movies were collected at a nominal magnification of 130,000x (physical pixel size: 0.463 Å/pixel) on a Titan Krios G4 (Thermo Fisher Scientific) operated at 300 kV and equipped with a Falcon 4i direct electron detector and Selectris energy filter set to a slit width of 10 eV. A defocus range of −0.8 to −1.8 um was used, with a total exposure time of 5 seconds fractionated into 100 0.05-second subframes. The total dose for each movie was 45 electrons/Å^2^. Movies were motion corrected using MotionCor2(52) in Scipion(53) and were Fourier cropped by a factor of 2 to a final pixel size of 0.927 Å/pixel. Motion corrected and dose-weighted micrographs were imported into RELION 5(54) for further image processing.

### Cryo-EM image processing

#### Mini-AD

Once imported into RELION 5 (54), the contrast transfer function of each micrograph was estimated using CTFFIND-4.1 (55). We used a modified version of Topaz for automated picking (56, 57). For training the Topaz neural network, we manually picked 1,605 filaments from 146 micrographs. From the resulting start-end coordinate pairs, 82,026 individual particle images were extracted, with a box size of 900 downscaled to 300 pixels and an inter-particle distance of 4.7Å. The coordinates of the individual particles were used to train the Topaz neural network and the resulting model was used to pick 3,858,586 particles from 7,411 micrographs. Multiple rounds of reference-free 2D classification were performed to separate different filament morphologies. Segments belonging the most abundant class, as described above, were re-extracted with a box size of 288 pixels without downscaling. RELION’s relion_helix_inimodel2d was used to generate an initial 3D volume. Multiple 3D classification jobs were run in an attempt to separate the best particles for 3D reconstruction, which resulted in the map used here.

#### dGAE seeded with mini-AD

Once imported into RELION 5 (54), the contrast transfer function of each micrograph was estimated using CTFFIND-4.1 (55).We used a modified version of Topaz for automated picking (56, 57). For training the Topaz neural network, we manually picked 592 filaments from 149 micrographs. From the resulting start-end coordinate pairs, 38,028 individual particle images were extracted, with a box size of 810 downscaled to 270 pixels and an inter-particle distance of 4.7Å. The coordinates of the individual particles were used to train the Topaz neural network and the resulting model was used to pick 1,627,798 particles from 4,290 micrographs. Multiple rounds of reference-free 2D classification were performed to separate different filament morphologies.

### Spin labeling of dGAE tau

Site-directed mutagenesis was used to introduce cysteine residues at positions Q351 and T373 in the dGAE construct, while the native cysteine at position 322 was mutated to serine to prevent non-specific labeling. All constructs were expressed and purified as described above. Spin labeling was carried out using (1-acetoxy-2,2,5,5-tetramethyl-Δ³-pyrroline-3-methyl) methanethiosulfonate (MTSL), obtained from Toronto Research Chemicals. Prior to labeling, proteins were treated with a 10-fold molar excess of dithiothreitol (DTT) to reduce cysteine residues, and excess DTT was removed through buffer exchange using a PD-10 desalting column (GE Healthcare). The reduced proteins were then incubated overnight at 4 °C with a 10-fold molar excess of MTSL per cysteine. Unreacted spin label was removed using a second PD-10 column. Labeling efficiency, determined by the molar ratio of attached spin labels to available cysteine residues, was approximately 90% for single-cysteine variants and 50–60% for double-cysteine constructs.

### Cw-EPR

CW-EPR measurements were performed at room temperature using a Bruker EMX X-band spectrometer operating at 9.8 GHz (EMX; Bruker Biospin, Billerica, MA) equipped with a dielectric resonator cavity (ER 4123D; Bruker Biospin, Billerica, MA). Mini-AD fibrils were prepared under conditions similar to those described above, using a mixture of 5% spin-labeled monomers (either end-to-end labeled or T377C) and 95% unlabeled monomers. A total of 1.6 mL of 100 μM monomer solution was incubated with heparin to induce fibrillization. Following fibril formation, samples were washed to remove unincorporated monomers using Amicon Ultra-0.5 (100 kDA cutoff) as previously described and concentrated to a final volume of 40 μL.

For cw-EPR, 5.0 μL of the fibril sample was loaded into a quartz capillary tube (0.6 mm internal diameter, CV6084; VitroCom), sealed at one end with Critoseal. Spectra were acquired using the following parameters: 6 mW microwave power, 0.5 G modulation amplitude, 100 G sweep width, and 30 signal-averaged scans.

The cw-EPR spectral component analysis was carried out using MultiComponent, a software package developed by Christian Altenbach and written in LabVIEW (National Instruments), available at: [UCLA Hubbell Lab Website](https://www.biochemistry.ucla.edu/Faculty/Hubbell/software.html) Fitting procedures followed previously described protocols for tau fibrils (31). Distance information was extracted using the software package ShortDistances (developed by Christian Altenbach using LabVIEW., available at: [UCLA Hubbell Lab Website](https://www.biochemistry.ucla.edu/Faculty/Hubbell/software.html)), following previously established protocols (32). See SI for more details on analysis.

### Double Electron Electron Resonance (DEER) of dGAE fibrils

During the fibrillization reaction, 5% of the spin-labeled dGAE (at sites Q351 and T373) were mixed with 95% non-spin-labeled dGAE to ensure that the spin-labeled monomers in a fibril were well separated from each other. This approach minimizes intermolecular spin-spin interactions, ensuring that only the desired distances are measured, as statistically, the distances falling within the DEER range should come from the intramolecular spin pairs, corresponding to the structure of individual dGAE monomers. Due to the random incorporation of spin-labeled monomers into the fibrils, any residual intermolecular interactions are also randomized, ensuring that such interactions do not produce unwanted distance distributions. These random interactions contribute to the background of the DEER signal, which is corrected during analysis.

For both mini-AD seeding dGAE and dGAE seeding dGAE reactions, the same conditions described above were used with minor modifications. Specifically, DTT was omitted from the fibrillization reactions involving spin-labeled proteins to prevent disruption of the spin labels. This omission had no observable impact on fibril formation. Fibrils formed with 5% spin-labeled monomers showed ThT fluorescence and negative-stain TEM profiles comparable to those of unlabeled fibrils. The dGAE seeds prepared without DTT were used for DEER experiments involving dGAE seeding dGAE.

To obtain sufficient material for DEER, the reactions were scaled up to a total of 500 μg of monomer. Following fibrillization, unreacted monomers were removed by overnight dialysis (molecular weight cut off 25 kDa) against ultrapure water (Invitrogen), and the fibrils were subsequently lyophilized. The dried fibrils were resuspended in 28 μL of D₂O-based 10 mM phosphate buffer and mixed with 12 μL of D₈-glycerol (30% v/v). The final mixture was transferred to a quartz tube with 2 mm internal and 3 mm external diameter and flash-frozen in liquid nitrogen for EPR measurements.

The DEER experiments were performed with a pulsed Q-band Bruker E580 Elexsys spectrometer, equipped with a Bruker QT-II resonator and a 300 W TWT amplifier with an output power of 20 mW for the recorded data (Applied Systems Engineering, Model 177Ka). The temperature of the cavity was maintained at 65 K using a Bruker/ColdEdge FlexLine Cryostat (Model ER 4118HV-CF100). The bridge is equipped with an Arbitrary Wave Generator to create shaped pulses for increased sensitivity.

The following DEER pulse sequence was used: π_obs_/2– τ_1_ – π_obs_ – (t-π_pump_) – (τ_2_-t) – π_obs_ – τ_2_ – echo. *V*(*t*) is recorded as the integral of the refocused echo as a function of time delay, t, between the Hahn echo and pump pulse. Rectangular observe pulses were used with lengths set to π_obs_/2 = 10-12 ns and π_obs_ = 20-24 ns, π_pump_ = 100 ns. The chirp pump pulse was applied with a frequency width of 60 MHz to excite a distinct spin population, referred to as B spins, while the observe pulse was set 33 G up field from the center of the pump frequency range to probe another distinct spin population, A spins *τ*_1_ was set to 180 ns and *τ*_2_ was set according to the SNR profile of the dipolar signal. Deuterium ESEEM suppression was done by incrementing *τ*_1_ with 16 ns steps, n=8 times. The data was acquired with resolution of 16 ns, 16-step phase cycling, and signal averaged until desirable SNR. The DEER data analysis was carried out with two methods, using DD-Gaussian and LongDistances (developed by Christian Altenbach using LabVIEW, Available at: [UCLA Hubbell Lab Website](https://www.biochemistry.ucla.edu/Faculty/Hubbell/software.html)) software. See SI appendix and Figure S7-S10 for further details about processing.

### Cell culture and stable line generation (cellular seeding studies)

A plasmid encoding mRuby3-Tau187(3R)-P2A-mClover3-Tau187(4R)-P301L was previously constructed and transfected into H4 neuroglioma cells using Lipofectamine 3000 (Thermo Fisher). Stable cell lines expressing the construct were selected by culturing in 500 μg/mL geneticin (G418), followed by fluorescence-activated cell sorting (FACS) using a Sony MA900 sorter to isolate populations co-expressing red (mRuby3) and green (mClover3) fluorescence.

Cells were maintained in Dulbecco’s Modified Eagle Medium (DMEM) supplemented with 10% fetal bovine serum (FBS) and 1% penicillin-streptomycin at 37°C in a humidified incubator with 5% CO₂. Cells were routinely passaged using 0.05% trypsin-EDTA upon reaching confluence.

### Preparation of sarkosyl-insoluble fractions (cellular seeding studies)

Brain tissue samples were weighed and homogenized in a Dounce homogenizer using 10 volumes of a freshly prepared homogenization buffer containing 0.8M NaCl, 1mM EGTA, 10% sucrose, 0.01M Na_2_H_2_P_2_O_7_, 0.1M NaF, 2mM Na_3_VO_4_, 0.025M β-glycerolphosphate, and 0.01M Tris–HCl at pH 7.4, supplemented with protease inhibitors (Roche, Switzerland). The homogenate was subjected to centrifugation at 20000 rcf for 22 minutes at 4° C, and the supernatant (referred to as SN1) was retained. The pellet was then resuspended in 5 volumes of the same buffer and centrifuged again at 20000 rcf for 22 minutes at 4° C. The second supernatant (SN2) was combined with SN1.

To this pooled supernatant (SN1 + SN2), 0.1% N-lauroyl sarcosinate (sarkosyl; Sigma-Aldrich) was added, and the mixture was agitated on a rotating shaker for one hour at room temperature. The solution was then centrifuged at 125,000 rcf for 63 minutes at 4 °C. After centrifugation, the supernatant was discarded, and the pellet, representing the sarkosyl-insoluble fraction, was rinsed and resuspended in 50mM Tris–HCl (pH 7.4) at a concentration of 200 µl per gram of initial tissue. Aliquots of 100 µl were stored at −80 °C for subsequent use.

### Fibril treatment (cellular seeding studies)

For fibril uptake experiments, cells were seeded at a density of 25,000 cells per well onto 96-well glass-bottom plates (CellVis) and allowed to adhere overnight. The following day, various fibrils were introduced using Lipofectamine 2000 (Invitrogen). Fibril complexes were prepared by mixing: 1.25 μL Lipofectamine 2000 with 8.75 μL Opti-MEM (Gibco, followed by the addition of 10 μL of fibril solution in Opti-MEM. The fibril-liposome complexes were incubated for 15 minutes at room temperature before being added to the cells. For peptide-based fibrils, a final concentration of 2 μM was used. For patient-derived Alzheimer’s disease (AD) fibrils, the concentration was empirically determined based on the formation of puncta in cells.

After 24 hours of fibril exposure, cells were fixed with 4% paraformaldehyde (PFA) in PBS for 10 minutes at room temperature. Fixed samples were washed three times with PBS and imaged using a Leica SP8 confocal fluorescence microscope with identical acquisition settings across conditions.

## Competing Interest Statement

K.S.K. consults for ADRx and Expansion Therapeutics and is a member of the Tau Consortium Board of Directors. V.V., S.L., A.L., K.T., K.K., and S.H. have filed for a patent based on the design of tau peptides presented in this paper. Patent information: Disc-ID-24-06-12-001.

## Author Contributions

V.V., M.V., A.P.L., S.H. designed research. S.L. designed simulations, V.V., G.E.M., S.L., A.P.L., K.T., A.Q., K.L.N., performed research. V.V., G.E.M., S.L., A.P.L., K.T., A.Q. analyzed data, A.A.M., E.T. helped with cryo-EM data collection, M.S.S., J.E.S., K.K., D.R.S., S.H. supervised research, V.V. and S.H. wrote the paper.

## Supporting information

Supporting information is attached as a separate file.

## Acknowledgements

The study of the role of disease mutation and selecting tauopathy-specific pathways was supported by the National Institutes of Health (NIH) under Grant Number R01AG05605. The study of the minimal prion design and seeding of tau was supported by the Tau Consortium of the Rainwater charitable fund. The cryo-EM sample preparation was supported by NIH MIRA under Grant Number R35GM136411. The W. M. Keck Foundation (www.wmkeck.org) supported the ongoing experimental and computational method developments for the tau shape propagation study.

Peptide Synthesis was performed at the Peptide Synthesis Core Facility of the Center for Regenerative Nanomedicine at Northwestern University. This facility has current support from the Soft and Hybrid Nanotechnology Experimental (SHyNE) Resource (NSF ECCS-2025633). The Center for Regenerative Nanomedicine, Northwestern University Office for Research, U.S. Army Research Office, and the U.S. Army Medical Research and Materiel Command have also provided funding to develop this facility. We thank Mark R. Karver, Director, CRN Peptide Synthesis Core for peptide synthesis.

We Acknowledge IMSERC (RRID:SCR_017874) at Northwestern University for EPR measurements.

The ns-TEM measurements in this work made use of the BioCryo facility (RRID:SCR_021288) of Northwestern University’s NUANCE Center, which has received support from the SHyNE Resource (NSF ECCS-2025633), the IIN, and Northwestern’s MRSEC program (NSF DMR-2308691).

We acknowledge staff and instrumentation support from the Structural Biology Facility at Northwestern University, the Robert H Lurie Comprehensive Cancer Center of Northwestern University and NCI CCSG P30 CA060553.

We acknowledge the use of the NRI-MCDB Microscopy Facility and the Resonant Scanning Confocal supported by NSF MRI grant DBI-1625770. The authors acknowledge the use of the Biological Nanostructures Laboratory within the California NanoSystems Institute, supported by the University of California, Santa Barbara and the University of California, Office of the President.

