## Supplementary Information for "Structure-specific Mini-Prion Model for Alzheimer’s Disease Tau Fibrils"

1. Department of Chemistry and Biochemistry, University of California, Santa Barbara, Santa Barbara, CA, USA. 2. Institute for Neurodegenerative Diseases, University of California San Francisco, San Francisco, CA, USA. 3. Department of Neurology, University of California San Francisco, San Francisco, CA, USA. 4. Neuroscience Research Institute, University of California Santa Barbara, Santa Barbara, CA, USA. 5. Department of Molecular, Cell and Developmental Biology, University of California Santa Barbara, Santa Barbara, CA, USA. 6. Department of Chemical Engineering, University of California, Santa Barbara, Santa Barbara, CA, USA. 7. Department of Biological Sciences, Northwestern University, Evanston, IL, USA. 8. Department of Chemistry, Northwestern University, Evanston, IL, USA. 9. Department of Physics, University of California, Santa Barbara, Santa Barbara, CA, USA. 10. Department of Biochemistry and Biophysics, University of California San Francisco, San Francisco, CA, USA.

*Corresponding author.

**Competing Interest Statement**: K.S.K. consults for ADRx and Expansion Therapeutics and is a member of the Tau Consortium Board of Directors. V.V., S.L., A.L., K.T., K.K., and S.H. have filed for a patent based on the design of tau peptides presented in this paper. Patent information: Disc-ID-24-06-12-001.

**Author Contributions:** V.V, M.V., A.P.L., S.H. designed research. S.L. designed simulations. V.V., G.E.M., S.L., A.P.L., K.T., A.Q., K.L.N., performed research. V.V., G.E.M., S.L., A.P.L., K.T., A.Q. analyzed data, A.A.M., E.T. helped with cryo-EM data collection, M.S.S., J.E.S., K.K., D.R.S., S.H. supervised research, V.V. and S.H. wrote the paper.

**This PDF file includes:**

Supporting text
Figures S1 to S11
SI References

### Supporting Text

#### Cw-EPR spectral component analysis and distance prediction

Cw-EPR spectral fitting was performed using MultiComponent (1, 2), a software package developed by Christian Altenbach and written in LabVIEW (National Instruments), available at: UCLA Hubbell Lab Website. Fitting procedures followed previously described protocols for tau fibrils (3). The mobile component was first determined by fitting the A, G, and R tensors to spectra of soluble monomeric peptides. The A and G tensors were optimized initially, followed by refinement of the rotational correlation time (R). A second component was then fit to fibrils formed with 5% spin labelled mini-AD with a single spin label at site T377C. These fibrils, while expected to exhibit minimal dipolar interaction and spin exchange, fall into the slow-motion regime due to their large size. The fibrils were previously characterized by an axially symmetric diffusion tensor ($\alpha_D$ = 0°, $\beta_D$ = 36°, $\gamma_D$ = 0°) and an ordering parameter of 20. For this component, only R was allowed to vary. A third component was added to model doubly labeled peptides incorporated into fibrils. This component shared the same R value as the second, reflecting similar dynamic regimes for all spin labels within the fibril. Additionally, a Heisenberg spin-exchange term ($\omega_{ss}$ = 140 MHz), empirically determined for tau fibrils, was incorporated into this component.

Distance information was extracted using the software package ShortDistances (1, 2), following previously established methods (4). Two CW-EPR spectra were collected for fibrils containing 5% spin-labeled protein: one from mini-AD fibrils labeled at residue T377, and another from end-to-end doubly labeled fibrils. Each spectrum was acquired with 2048 points and converted to single-column ASCII format. The spectrum from the doubly labeled fibrils was imported as the dipolar broadened spectrum (D), while the spectrum from T377-labeled fibrils served as the reference for the singly labeled (S), non-interacting condition. The dipolar spectrum was then fit to a Gaussian distance distribution, allowing the center distance, width, percentage of spin pairs contributing to the dipolar interaction, and the percentage of non-interacting spins to vary as fitting parameters.

#### DEER data Processing Workflow

This section contains the detailed procedure and program link for the DEER data processing present in this paper.

- ***DD Gaussian***

Raw $V(t)$ from Bruker Elexsys data file was analyzed using DD (5, 6) version 7C software package for MATLAB. Using the DD MATLAB GUI, the time traces were phase corrected, zero-time corrected, and the end of the signal was eliminated by 100-200 ns to eliminate artifact. The background fit was performed using the "Exponential" background option, while allowing the modulation depth, $\lambda$, concentration, and dimensionality, $d$, to vary. Once background fit was done with the 1-gaussian model, the background parameters were frozen to limit fitting parameters. The background corrected data was fitted with various number of gaussians, where the number of gaussian with the lowest reduced chi square and/or with the lowest Bayesian criteria (BIC) was chosen. The uncertainty analysis was estimated from the variance-covariance matrix method. The distance distribution fits are presented with 95% confidence interval.

To estimate the relative population of each expected structure, the area under the mean P(R) distribution corresponding to the predicted distance range for each structure was calculated. In cases where the distributions overlapped, the shared area was included in the population analysis of both structures. The total population was then normalized to 1.

- ***LongDistances***

Raw $V(t)$ from Bruker Elexsys data file was loaded into Long Distances (1, 2) for data analysis. For all samples, the background was fitted with dimensionality, $b_3 = 2$ to best approximate spin labels distributed across protofilament, using the VariableD model: $B(t) = b_1 * e^{[(b_2 t)^{\frac{b_3}{3}}]}$. Prior to background correction, the time traces were phase-corrected and the end of the signal was trimmed to eliminate experimental artifacts. The L curve was used as a criterion for determining the optimum smoothness parameter for the Model Free analysis by Tikhonov regularization. The uncertainty analysis was done using bootstrapping method with 100 samples. The distance distribution fits are presented with ±2 standard deviations (equivalent to 95% confidence interval).

**Alphafold Simulations for structure prediction**

AlphaFold2 (AF2) (7) structures were predicted using the ColabFold implementation (github.com/YoshitakaMo/localcolabfold) of AlphaFold2 using all default settings. The AF2 input was five copies of the mini-AD sequence, with the residue index increasing by 20 in between subsequent copies (i.e. putting 20 "U"s between each "VQIVYKPGGGNHKLTF"). The output structure was not sensitive to the residue index increase between copies. Note: we use the default single-chain model (alphafold2_ptm) rather than the AF2-multimer model for predictions; AF2-multimer does not appear to produce amyloid structures with cross β-sheet structure and side chain zippers. The template model is not used.

AlphaFold3 (AF3) structures were predicted using the alphafoldserver.com. Five copies of the mini-AD sequence were input with templates turned off and seed = 1.

The simulated density map of the predicted AF2 structure with three mini-AD folds was generated in UCSF ChimeraX (8) using the *molmap* command at a resolution of 2.4 Å.

### Figures

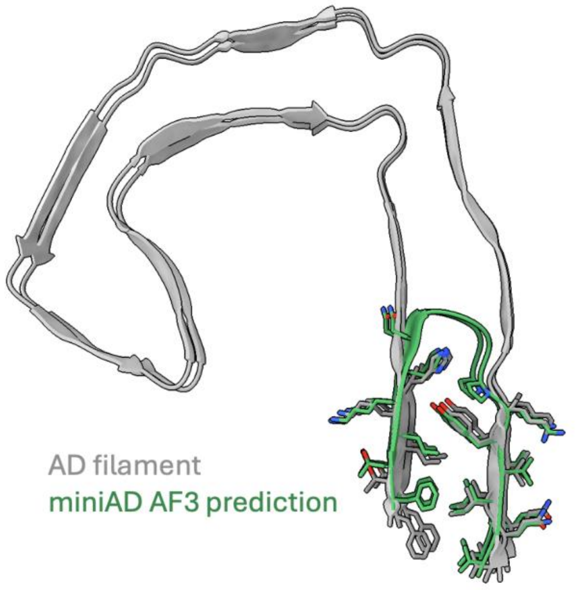


**Figure S1:** Mini-AD AF3 prediction fragment shown alongside the cryo-EM structure of the AD filament (PDB #5O3L). The AF3 prediction was made with 5 copies of the miniAD sequence, of which two are shown. The alpha carbon RMSD of resid 306-311 & 374-378 are 0.08 nm between the two dimer structures. The AF2 prediction also matched the cryoEM structure with a dimer alpha carbon RMSD of 0.48 nm. (See above for further details of the calculations).

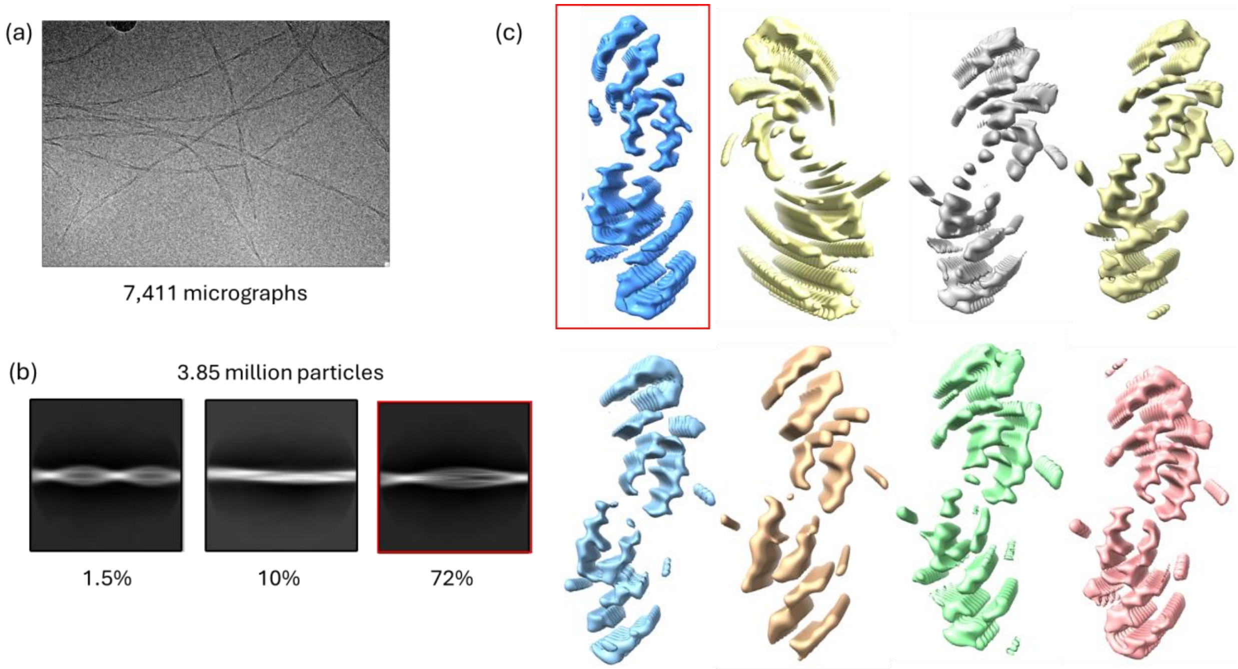


**Figure S2:** (a) Representative micrograph of mini-AD filaments. (b) Representative 2D class averages of the filaments from (a). The red box indicates the morphology which was selected for further processing. (c) Results from 3D classification, showing cross-sectional views of the filaments. The overall shape of each subunit closely resembles the designed peptide, assembling in a dimer-of-dimers arrangement, where two U-shaped mini-AD folds form a dimer that further associates with a second dimer to create a tetrameric structure.

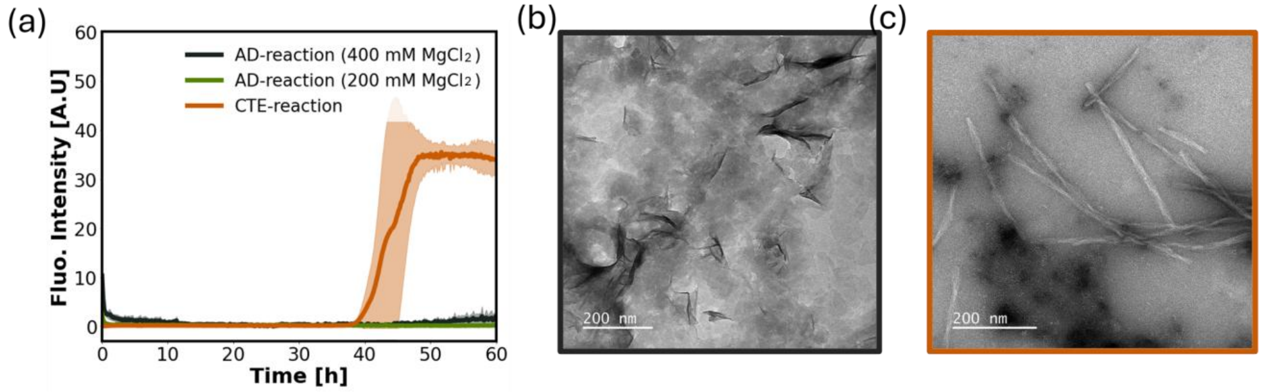


**Figure S3:** (a) Thioflavin T (ThT) fluorescence kinetics for unseeded AD and CTE reactions. The brick-colored curve represents the CTE reaction, while the olive green and black curves represent AD reaction conditions with 200 mM and 400 mM $MgCl_2$, respectively. Baseline correction was performed by subtracting the minimum fluorescence intensity from each curve. (b, c) Representative negative-stain TEM images taken after 60 hours of incubation for (b) the AD reaction (400 mM $MgCl_2$) and (c) the CTE reaction, the latter showing heterogeneous fibril populations. Scalebars represent 200 nm for each image.

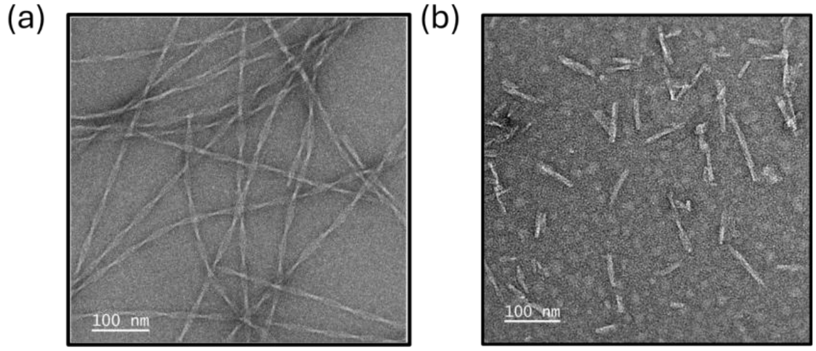


**Figure S4:** Representative negative-stain TEM images of mini-AD fibrils (a) before and (b) after sonication, with the latter showing fragmented fibrils ranging from 50 to 100 nm in length. Scalebars represent 100 nm.

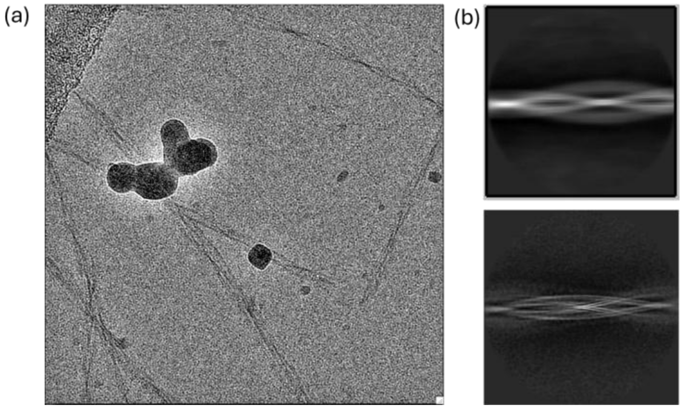


**Figure S5:** (a) Representative micrograph from the mini-AD seeded dGAE filaments. b) Representative 2D class averages of the filaments from (a) (top) and with a published 2D class average of CTE filaments (bottom) (9) for comparison.

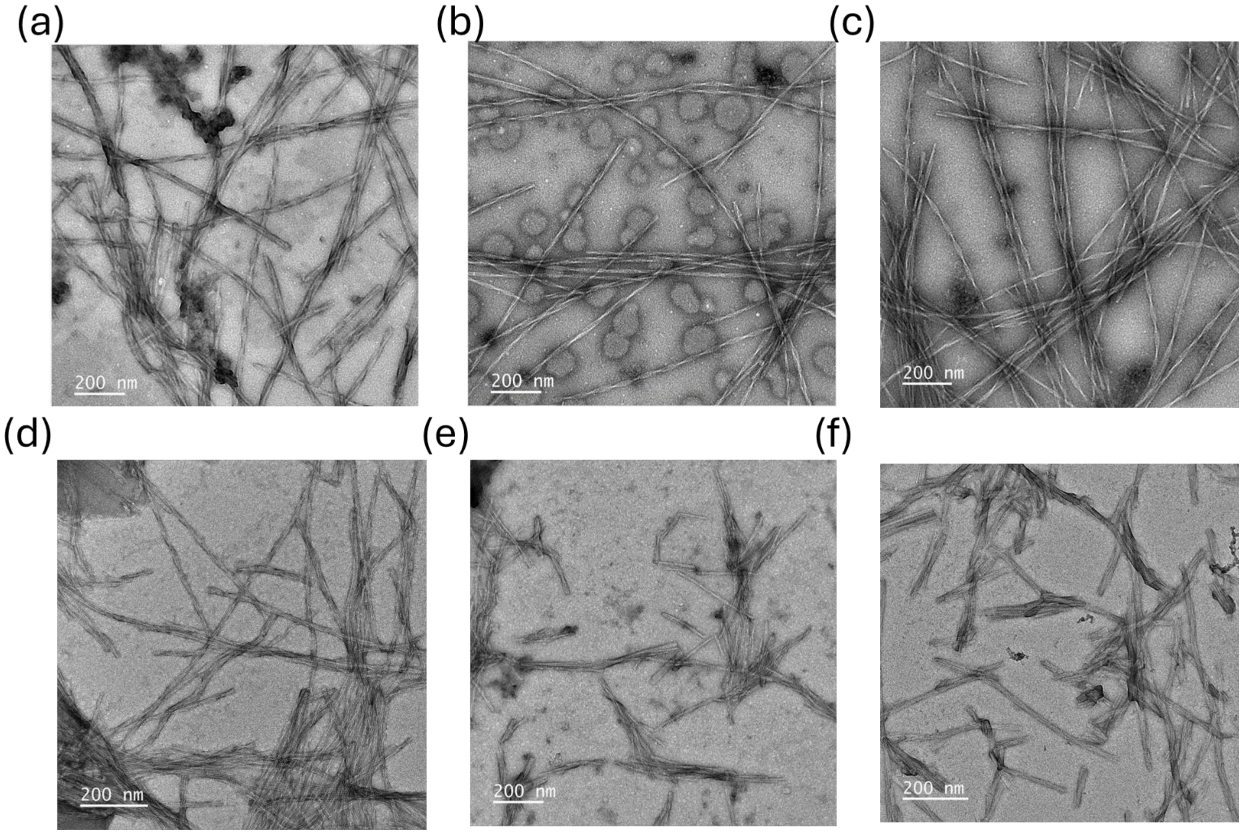


**Figure S6:** Representative negative-stain TEM images of dGAE fibrils generated through multiple rounds of seeding under different reaction conditions. (a–c) AD-reaction condition: (a) AD-dGAE$_{ms}$, (b) second-generation Gen2-AD-dGAE, and (c) third-generation Gen3-AD-dGAE showing progressive amplification of long, slender fibrils with PHF-like morphology. (d–f) CTE-reaction condition: (d) CTE-dGAE$_{ms}$, (e) Gen2-CTE-dGAE, and (f) Gen3-CTE-dGAE exhibiting thicker, less twisted structures that remain largely consistent across generations.

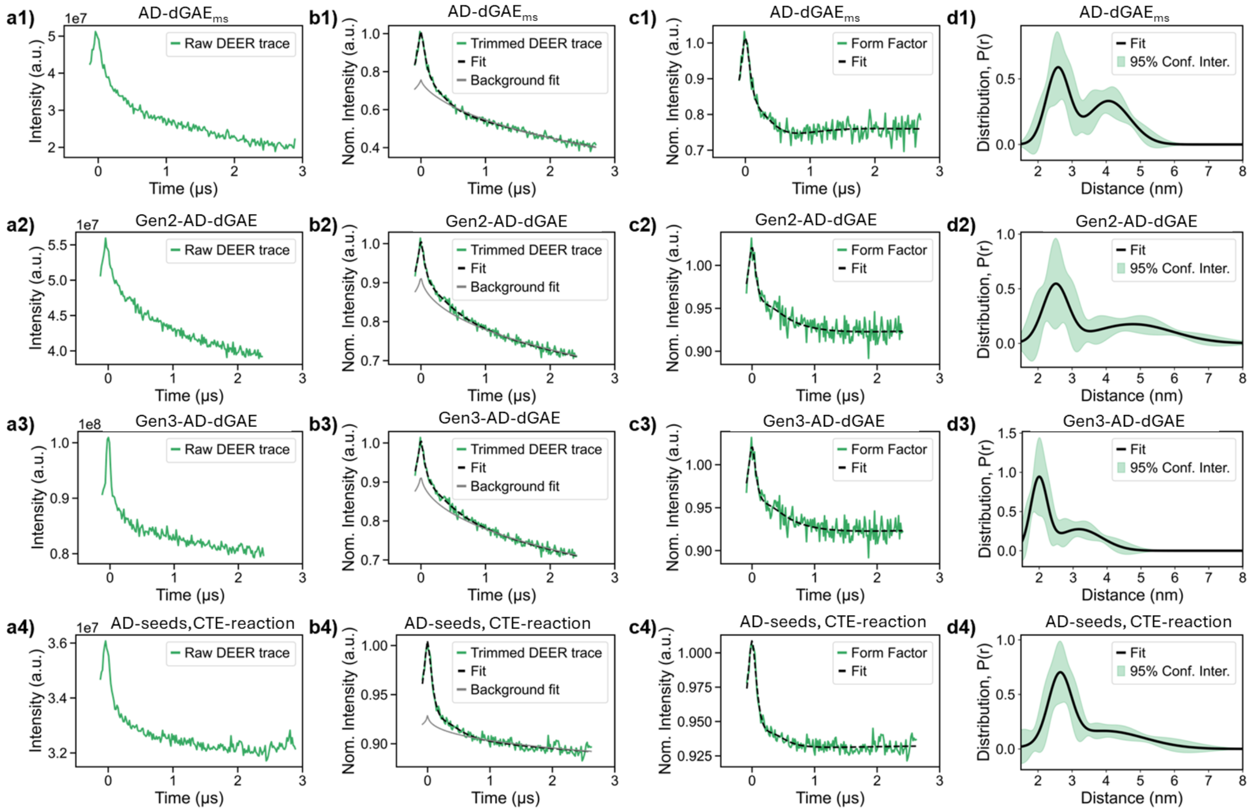


**Figure S7:** AD seeding series DEER data processed with DD using Gaussian Model. Row 1 (a1, b1, c1, d1) represents mini-AD seeded DGAE in AD-reaction (AD-dGAE$_{ms}$). Row 2 (a2, b2, c2, d2) represents Gen2-AD-dGAE. Row 3 (a3, b3, c3, d3) represents Gen3-AD-dGAE. Row 4 (a4, b4, c4, d4) represents dGAE fibrils seeded with AD-dGAE$_{ms}$ in CTE-reaction conditions. Column 1 (a1-a4) represents the raw DEER trace. Column 2 (b1-b4) represents the trimmed DEER trace, fit of the trimmed DEER trace, and background fit of the trimmed DEER trace. Column 3 (c1-c4) represents the form factor (trimmed DEER trace after background division) and its fit. Column 4 (d1-d4) represents the distance distribution with 95% confidence interval.

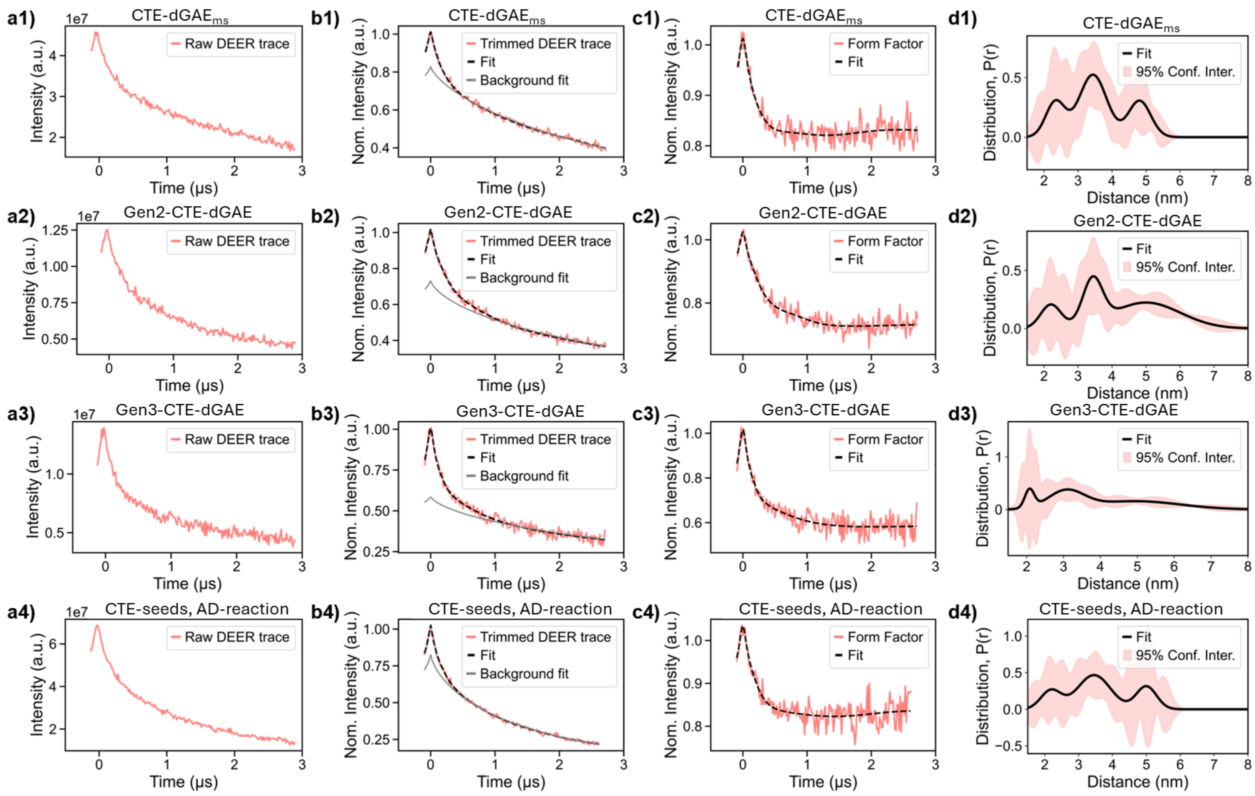


**Figure S8:** CTE seeding series DEER data processed with DD using Gaussian Model. Row 1 (a1, b1, c1, d1) represents mini-AD seeded DGAE in CTE-reaction (CTE-dGAE$_{ms}$). Row 2 (a2, b2, c2, d2) represents Gen2-CTE-dGAE. Row 3 (a3, b3, c3, d3) represents Gen3-CTE-dGAE. Row 4 (a4, b4, c4, d4) represents dGAE fibrils seeded with CTE-dGAE$_{ms}$ in AD-reaction conditions. Column 1 (a1-a4) represents the raw DEER trace. Column 2 (b1-b4) represents the trimmed DEER trace, fit of the trimmed DEER trace, and background fit of the trimmed DEER trace. Column 3 (c1-c4) represents the form factor (trimmed DEER trace after background division) and its fit. Column 4 (d1-d4) represents the distance distribution with 95% confidence interval.

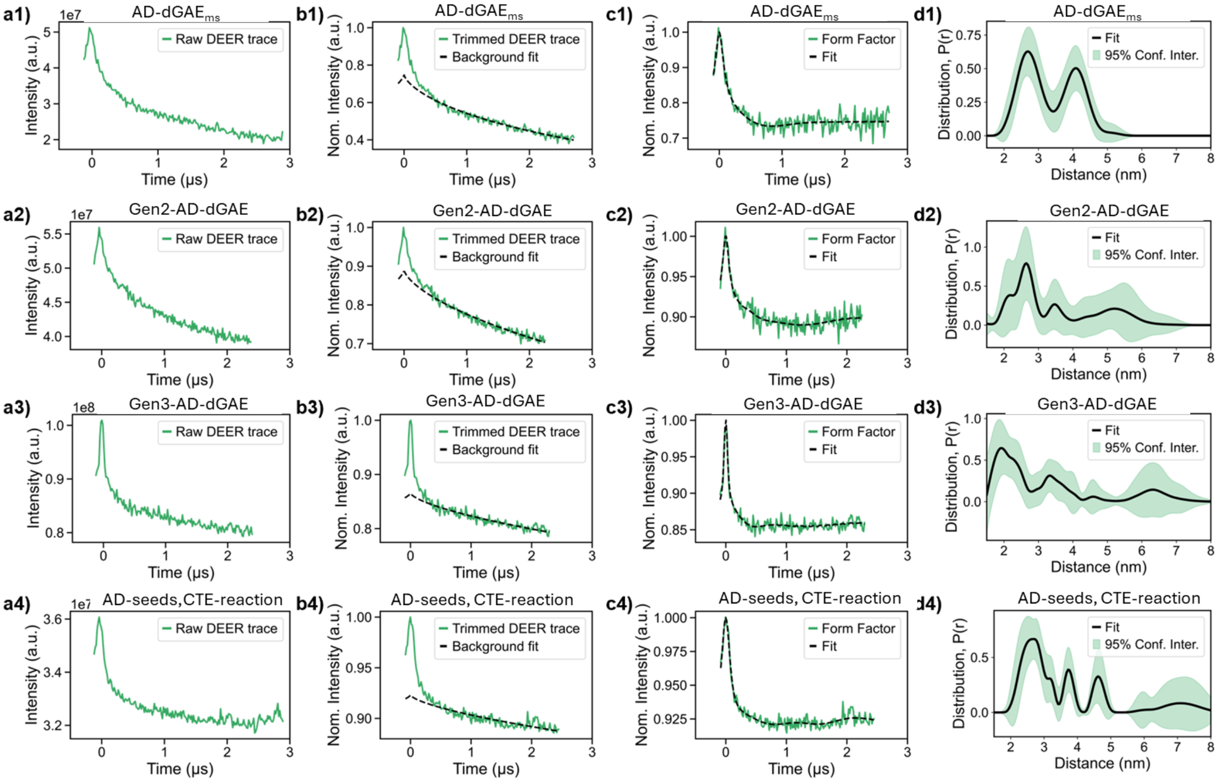


**Figure S9:** AD data processed with LongDistance software. Row 1 (a1, b1, c1, d1) represents mini-AD seeded DGAE in AD-reaction (AD-dGAE$_{ms}$). Row 2 (a2, b2, c2, d2) represents Gen2-AD-dGAE. Row 3 (a3, b3, c3, d3) represents Gen3-AD-dGAE. Row 4 (a4, b4, c4, d4) represents dGAE fibrils seeded with AD-dGAE$_{ms}$ in CTE-reaction conditions. Column 1 (a1-a4) represents the raw DEER trace. Column 2 (b1-b4) represents the trimmed DEER trace, fit of the trimmed DEER trace, and background fit of the trimmed DEER trace. Column 3 (c1-c4) represents the form factor (trimmed DEER trace after background division) and its fit. Column 4 (d1-d4) represents the distance distribution with 95% confidence interval.

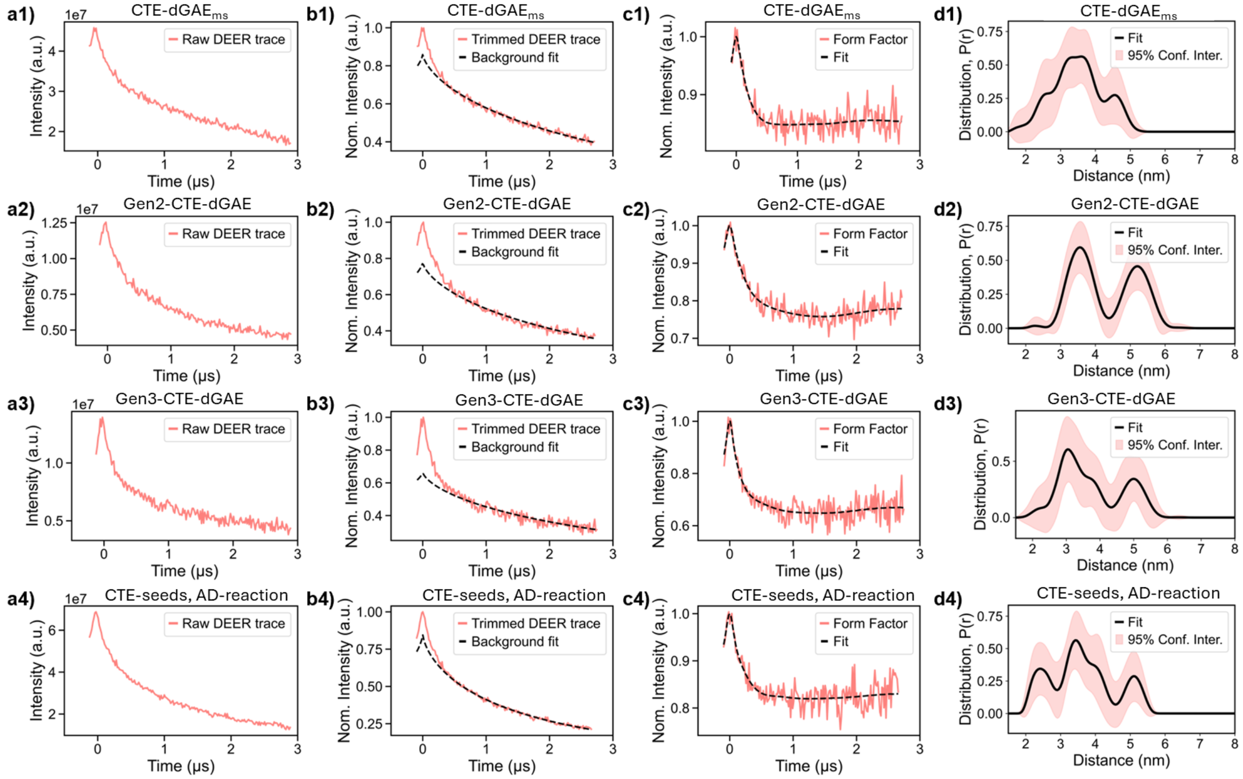


**Figure S10:** CTE data processed with LongDistance. Row 1 (a1, b1, c1, d1) represents mini-AD seeded DGAE in CTE-reaction (CTE-dGAE$_{ms}$). Row 2 (a2, b2, c2, d2) represents Gen2-CTE-dGAE. Row 3 (a3, b3, c3, d3) represents Gen3-CTE-dGAE. Row 4 (a4, b4, c4, d4) represents dGAE fibrils seeded with CTE-dGAE$_{ms}$ in AD-reaction conditions. Column 1 (a1-a4) represents the raw DEER trace. Column 2 (b1-b4) represents the trimmed DEER trace, fit of the trimmed DEER trace, and background fit of the trimmed DEER trace. Column 3 (c1-c4) represents the form factor (trimmed DEER trace after background division) and its fit. Column 4 (d1-d4) represents the distance distribution with 95% confidence interval.

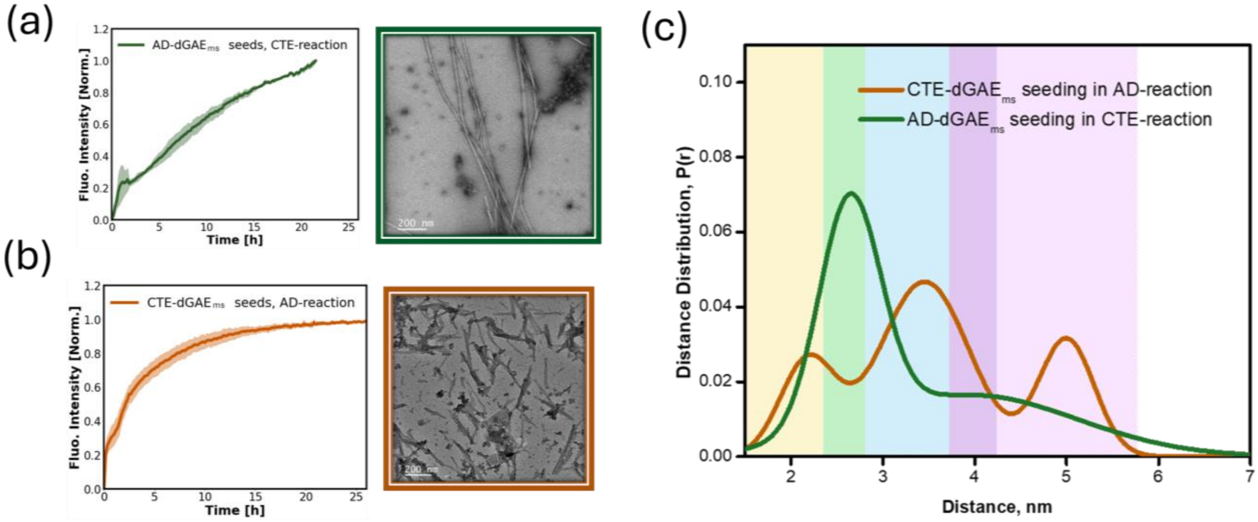


**Figure S11:** (a, b) Normalized ThT kinetics (left) and TEM images (right) of fibrils generated by cross-condition seeding: (a) AD-dGAE$_{ms}$ seeding dGAE under CTE-reaction conditions, and (b) CTE-dGAE$_{ms}$ seeding dGAE under AD-reaction conditions. (c) DEER P(r) distributions for fibrils from these cross-seeding reactions: AD-dGAE$_{ms}$ in CTE-reaction (dark green) and CTE-dGAE$_{ms}$ in AD-reaction (brown).
